# Two separation-of-function isoforms of human TPP1 and a novel intragenic noncoding RNA dictate telomerase regulation in somatic and germ cells

**DOI:** 10.1101/626523

**Authors:** Sherilyn Grill, Kamlesh Bisht, Valerie M. Tesmer, Christopher J. Sifuentes, Jayakrishnan Nandakumar

## Abstract

Telomerase replicates chromosome ends in germ and somatic stem cells to facilitate continued proliferation. Telomerase action depends on the telomeric protein TPP1, which recruits telomerase to telomeres and facilitates processive DNA synthesis. Here we identify separation-of-function long (TPP1-L) and short (TPP1-S) isoforms of TPP1 differing only in 86 amino acids at their N-terminus. While both isoforms retain the ability to recruit telomerase, only TPP1-S facilitates telomere synthesis. We identify a novel intragenic noncoding RNA in the 3’-UTR of the TPP1-encoding gene that specifically shuts down telomerase activation-incompatible TPP1-L to establish TPP1-S as the predominant isoform in somatic cells. Strikingly, TPP1-L is the major isoform in testes, where it can function to restrain telomerase in mature germ cells. Our studies uncover how differential expression of two isoforms allows TPP1 to perform separate functions in different cells, and demonstrate how isoform choice can be determined by an intragenic noncoding RNA.

## Introduction

The end replication problem arises due to incomplete chromosome end synthesis by DNA polymerases (Levy et al., 1992). This leads to the gradual loss of DNA at the ends of linear chromosomes during every replication cycle. This chromosome shortening sets a limit on the number of times most somatic cells can divide, thereby providing a natural anti-tumorigenic mechanism in large, long-lived mammals such as humans (Gomes et al., 2011). However somatic and germline stem cells must preserve their ability to self renew over long periods of time, lasting as long as the life of the individual itself. Telomerase, a unique enzymatic ribonucleoprotein (RNP) complex, is a reverse transcriptase that synthesizes DNA at the 3’ ends of chromosomes (Greider and Blackburn, 1985). Using a template sequence in its RNA subunit (TR) and a reverse transcriptase protein subunit (TERT), telomerase synthesizes multiple telomeric hexad repeats (GGTTAG in mammals) at chromosome ends, compensating for the attrition from incomplete DNA replication (Greider and Blackburn, 1989; Lingner et al., 1997; Meyerson et al., 1997). Not surprisingly, germline mutations in the core subunits of telomerase or in genes important for telomerase function result in stem cell dysfunction diseases that are collectively referred to as telomeropathies (Dokal, 2011; Savage, 2014).

While reduced telomerase function in stem cells can result in telomeropathies, aberrant reactivation of telomerase in somatic cells that do not normally express telomerase is a hallmark of an overwhelming majority of cancers (Kim et al., 1994). Thus telomerase must be tightly regulated, requiring sustained expression in stem cells, but complete shutdown upon differentiation. Interestingly, telomerase is constitutively expressed in most somatic cells of smaller short-lived mammals such as rodents (Gomes et al., 2011; Prowse and Greider, 1995). This presumably provides a proliferative advantage for these mammals, as fewer total cell divisions in their lifetime mitigate chances of accumulating oncogenic mutations.

Human telomeres are composed of several kilobases long telomeric DNA repeats bound to multiple copies of a six-member protein complex called shelterin (Fig. 1A) (Palm and de Lange, 2008). The primary function of shelterin is to protect natural chromosome ends from being recognized as double-stranded (ds) DNA breaks requiring repair. While TRF1 and TRF2 bind the ds fraction of telomeric DNA (Broccoli et al., 1997), POT1 binds with high specificity and affinity to the short (50-200 nucleotides) single-stranded (ss) 3’ overhang composed of GGTTAG repeats (Baumann and Cech, 2001; Lei et al., 2004). POT1 binds TPP1 to form a heterodimer with greater affinity for ss telomeric DNA than POT1 alone (Wang et al., 2007). The TIN2 protein connects TPP1 to TRF1 and TRF2 (Frescas and de Lange, 2014; Kim et al., 1999), while the sixth shelterin component, Rap1, constitutively binds TRF2 (Li et al., 2000).

**Figure 1.**
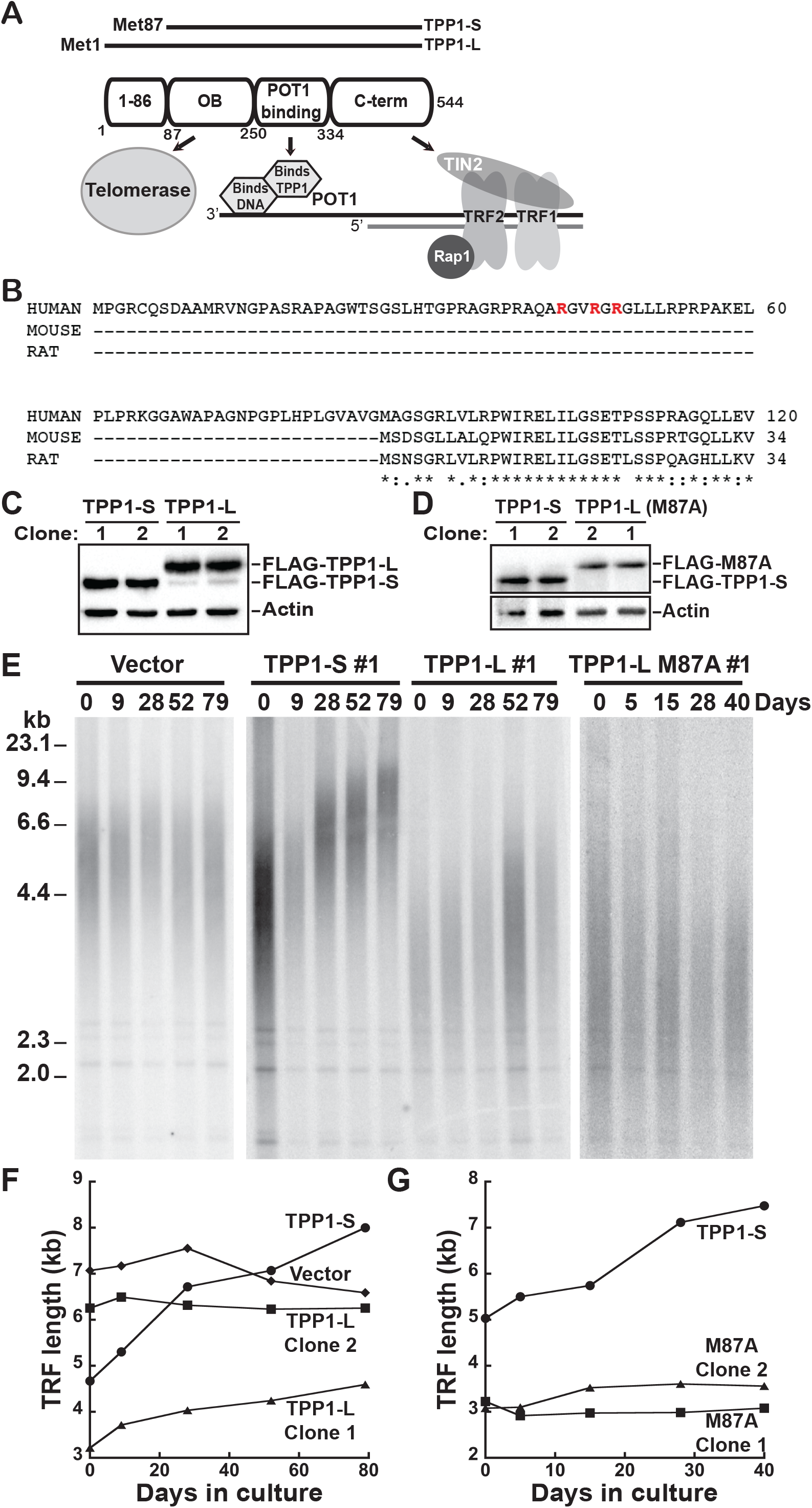
TPP1-S but not TPP1-L overexpression causes robust telomere elongation. (A) Schematic showing a TPP1-centric view of the shelterin complex in humans. Amino acid numbering below the domain diagram indicates domain boundaries. The different N-termini for TPP1-L and TPP1-S are indicated with horizontal lines above the domain diagram. (B) Primary sequence for the N-terminus of human, mouse, and rat TPP1 proteins. Asterisks indicate complete conservation, colons indicate sequence similarity between all three species, and dots indicate sequence similarity between two species only. Arginine residues in red indicate those mutated in the R_3_E_3_ mutant. (C) Immunoblot showing expression levels of FLAG-TPP1-S and FLAG-TPP1-L protein from indicated stable cell lines using anti-FLAG-HRP conjugate antibody. The actin immunoblot serves as a loading control. (D) Immunoblot of FLAG-TPP1-S and FLAG-TPP1-L M87A protein expression levels from indicated stable cell lines using anti-FLAG-HRP conjugated antibody. (E) Telomere restriction fragment (TRF) Southern blot analysis of cultured HeLa-EM2-11ht clonal cell lines stably expressing vector or the indicated TPP1 isoform. (F and G) Plot of the mean telomere restriction fragment (TRF) length for HeLa-EM2-11ht cells stably expressing vector or the indicated constructs as a function of days in culture. See also Figure S1.

Although shelterin is well suited for protecting chromosome ends, it also provides a mechanism for recruiting telomerase to its substrate at the very 3’ end of telomeres (Fig. 1A) (Nandakumar and Cech, 2013). This is achieved by an OB (<u>o</u>ligonucleotide/oligosaccharide-<u>b</u>inding) fold domain in the shelterin protein TPP1 (encoded by the *ACD* gene), which recruits telomerase to telomeres (Fig. 1A) (Abreu et al., 2010; Xin et al., 2007). Once recruited, telomerase synthesizes telomeric DNA with high processivity in a POT1-TPP1 dependent manner. Two regions in the OB domain, the TEL (<u>T</u>PP1’s glutamate (<u>E</u>) and leucine (<u>L</u>) rich) patch and the NOB (<u>N</u>-terminus of <u>OB</u> domain), are critical for all of TPP1’s telomerase-associated functions, including telomere lengthening in dividing human cells (Grill et al., 2018; Nakashima et al., 2013; Nandakumar et al., 2012; Sexton et al., 2012; Zhong et al., 2012). Indeed a naturally-occurring mutation, K170Δ, adjacent to two critical TEL patch residues has been shown in two separate individuals to result in telomeropathies (Bisht et al., 2016; Guo et al., 2014; Kocak et al., 2014). The NOB region is remarkable in that despite its critical importance it is not strictly conserved between mouse and human TPP1 homologs. Simply switching amino acids in the mouse TPP1 NOB with the equivalent human NOB residues allows mouse TPP1 to stimulate human telomerase, highlighting NOB as a key element for dictating mouse versus human species specificity of the TPP1-telomerase interaction (Grill et al., 2018).

A second distinct difference at the N-terminus between the human and mouse TPP1 orthologs was noted during their discovery. Human TPP1 was originally annotated to encompass 544 amino acids (aa) initiating at Met1 (Fig. 1A) (Houghtaling et al., 2004; Liu et al., 2004; Ye et al., 2004); we refer to this isoform as TPP1-L (and will maintain this numbering scheme). As such a majority of studies on human TPP1 involved TPP1-L overexpression (encoded by cDNA) (Abreu et al., 2010; Sexton et al., 2014; Tejera et al., 2010; Ye et al., 2004; Zhong et al., 2012). Yet, the existence of a shorter isoform that initiates at Met87, and referred to here as TPP1-S, was suggested by the realization that rodent TPP1 unequivocally initiates at a Met equivalent to human TPP1 Met87 (Fig. 1A,B) (Hockemeyer et al., 2006; Hockemeyer et al., 2007; Houghtaling et al., 2004; Nandakumar et al., 2012; Xin et al., 2007). Thus the TPP1-OB crystal structure (Wang et al., 2007) and other functional studies depicting the telomerase-related functions of TPP1 have used TPP1-S protein (Hwang et al., 2012; Nandakumar et al., 2012; Schmidt et al., 2014). Although TPP1-L is also able to recruit telomerase to the telomere (Abreu et al., 2010; Zhong et al., 2012), telomere hyperelongation that is characteristic of TPP1-S overexpression was not observed for TPP1-L (Houghtaling et al., 2004; Nandakumar et al., 2012; Xin et al., 2007; Ye et al., 2004).

Here we clarify the function and representation of TPP1-L and TPP1-S isoforms in human cells. We show that both TPP1-S and TPP1-L can recruit telomerase to the telomere, but only TPP1-S can activate telomere synthesis. We further show that TPP1-L is shut down by a noncoding RNA (ncRNA) expressed from its own 3’-UTR, revealing a novel, intragenic ncRNA-mediated mechanism for isoform-specific gene silencing. TPP1-S is the predominant isoform in almost all human cells, including human embryonic stem cells, cancer cells, and other somatic cells. Strikingly, TPP1-L expression is specifically upregulated during spermatogenesis such that TPP1-L is the predominant isoform of TPP1 in testes. We envision that TPP1-L prevents undesired telomere elongation during the non-mitotic stages of spermatogenesis, a unique biological niche where telomerase could persist despite a lack of active cell division. These results unveil a distinct separation-of-function between two isoforms of human TPP1 in telomerase activation, and discover a novel intragenic ncRNA that differentially regulates the isoforms to accomplish unique goals in both germline and somatic cells.

## Results

### TPP1-S, but not TPP1-L, overexpression causes hyperelongation of telomeres

To directly compare the effects of TPP1-L and TPP1-S in telomere length regulation, we used a single-site integration, doxycycline (dox)-inducible stable cell line strategy that we have used to characterize TEL patch and NOB mutants of TPP1-S (Grill et al., 2018; Nandakumar et al., 2012). We engineered a stable cell line that expresses C-terminally FLAG-tagged TPP1-L (aa 1-544) and compared it directly to a previously characterized C-terminally FLAG-tagged TPP1-S (aa 87-544) stable cell line. Individual clones were selected to equalize TPP1 protein level. In all TPP1-L clones we also observed a modest amount of shorter FLAG-tagged protein that was similar in size to that of FLAG-tagged TPP1-S (Fig. 1C). We speculated that the M87 start codon in TPP1-L cDNA was being utilized to produce FLAG-tagged TPP1-S protein, which could have complicated the interpretation of previous studies that used TPP1-L. To circumvent this potential caveat, we engineered another HeLa cell line that stably expresses FLAG-tagged TPP1-L protein harboring an M87A mutation. As aa 87-90 are absent in TPP1-S constructs that have been shown to fully sustain end protection and telomerase-stimulatory functions (Wang et al., 2007) (called TPP1-N in the literature; referred to here as TPP1^90-334^), we reasoned that the M87A mutation would be benign to TPP1-L’s functions. As expected, the M87A mutation blocked the production of the shorter TPP1 protein (Fig. 1D). To determine how TPP1-L and TPP1-L M87A regulate telomere length relative to TPP1-S, we performed telomere restriction fragment (TRF) length analysis of our stable cell lines (Fig. 1E and Fig. S1A). A stable cell line containing the dox-inducible promoter but an empty cassette (Vector) was assayed in parallel as a negative control. Consistent with previous results, overexpression of TPP1-S resulted in the characteristic lengthening of telomeres over time (+42 bp/day), while telomeres of the vector cell line remained relatively stable (−6 bp/day) (Fig. 1F; Fig. S1B) (Nandakumar et al., 2012; Xin et al., 2007). In contrast, overexpression of TPP1-L resulted in either no lengthening (0 bp/day) or a moderate lengthening (17 bp/day) of telomeres (Fig. S1B). The observed low level of telomere length maintenance in TPP1-L overexpressing cells appears to be due to co-expression of TPP1-S protein as the TPP1-L M87A line had short telomeres (~3 kb) that did not elongate over time (Fig. 1D,E,G; Fig. S1A,C). TPP1-S and TPP1-L appear to play contrasting roles in telomere length maintenance as TPP1-S, but not TPP1-L, enhances telomere elongation.

### Both TPP1-S and TPP1-L recruit telomerase to telomeres

Telomere shortening through overexpression of TPP1-S TEL patch or NOB mutants is caused by a decreased association of these proteins with telomerase (Grill et al., 2018; Nandakumar et al., 2012). We asked if TPP1-L M87A was impaired in its ability to recruit telomerase to telomeres. To evaluate this, we performed immunofluorescence (IF) and fluorescence *in situ* hybridization (FISH) to visualize the localization of FLAG-TPP1 and TR in HeLa nuclei. In agreement with our observed telomere lengthening (Fig. 1E,F) and previously published data, telomerase (TR) co-localized with over 90 percent of TPP1-S foci, indicative of robust telomerase recruitment to telomeres (Fig. 2A,B) (Nandakumar et al., 2012). We also observed robust recruitment of telomerase to telomeres in both TPP1-L M87A clones (91% and 84% for clone 1 and clone 2 respectively; Fig. 2A,B; Fig. S2A). Furthermore, this phenotype was recapitulated in both TPP1-L WT clones (Fig. S2B). These data further distinguish TPP1-L from the previously characterized TEL patch and NOB mutants of TPP1-S.

**Figure 2.**
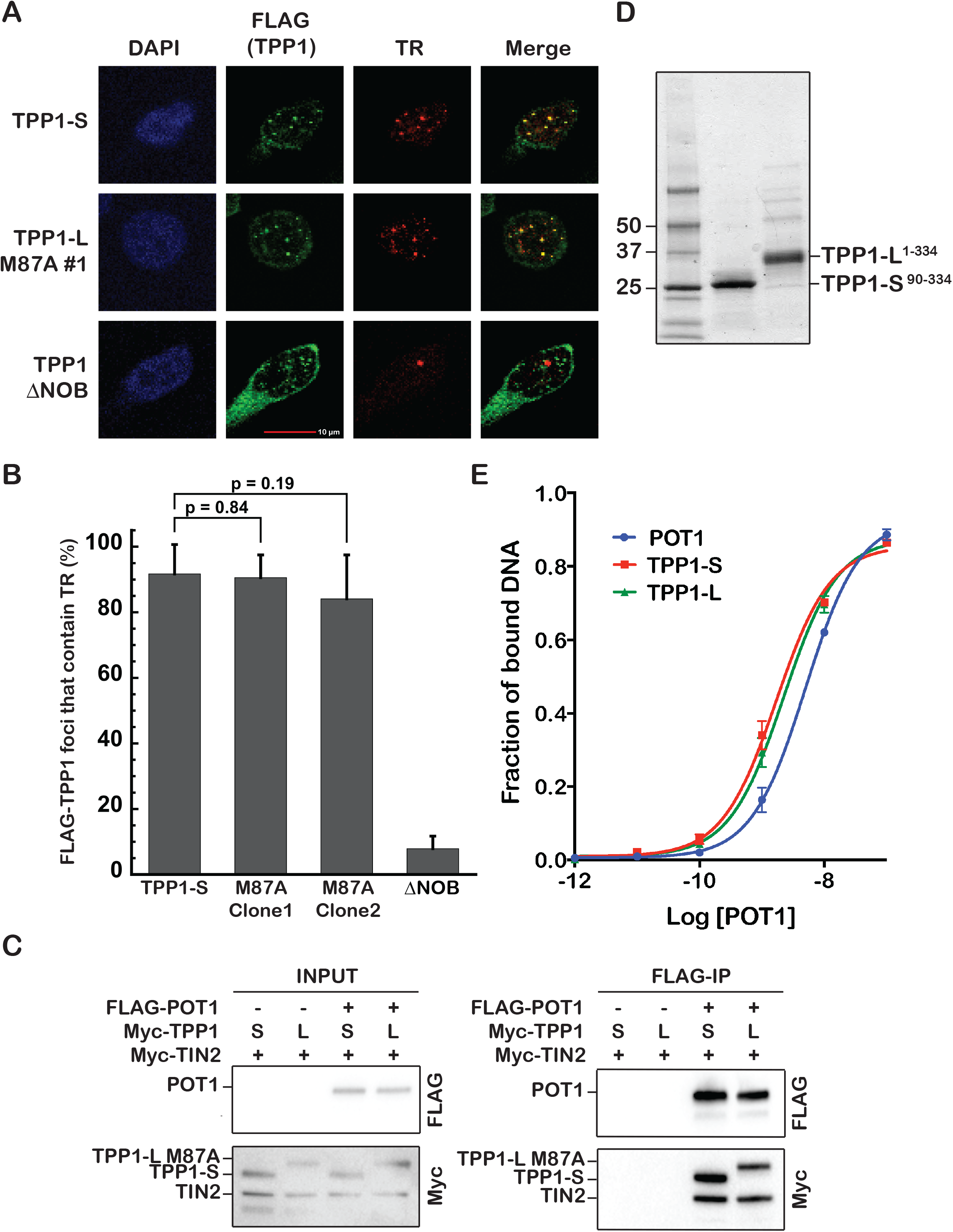
TPP1-S and TPP1-L recruit telomerase and protect chromosome ends. (A) HeLa-EM2-11ht cells expressing TPP1-S, TPP1-L M87A, or TPP1ΔNOB were analyzed for telomerase recruitment to telomeres by immunofluorescence and fluorescence *in situ* hybridization. “FLAG (TPP1)” indicates immunofluorescence signal of the indicated TPP1 construct at telomeres (green). Telomerase RNA (“TR”) was detected by fluorescence *in situ* hybridization with a fluorescently tagged DNA probe (red). The “Merge” panel depicts the extent of telomerase recruitment to telomeres (yellow). (B) Quantitation of telomerase recruitment data of which A is representative. For each clone >100 telomere foci were scored and the mean percentage (bar) and standard deviation (error bar) of FLAG-TPP1 foci containing TR was plotted for triplicate measurements. Statistical significance was scored with a two-tailed student’s t test. (C) Pull down of transiently expressed FLAG-POT1 on anti-FLAG conjugated beads with indicated Myc-TPP1 and Myc-TIN2 constructs. “INPUT” immunoblots refer to lysates before incubation with anti-FLAG beads while “FLAG-IP” immunoblots refer to anti-FLAG beads after immunoprecipitation and wash steps. (D) SDS-PAGE analysis of purified TPP1-S^90-334^ and TPP1-L^1-334^ proteins used in DNA-binding and telomerase primer extension experiments. (E) DNA binding curves from filter binding analysis of increasing concentrations of purified POT1 protein in the absence of any TPP1 protein, or in the presence of (200 nM) of either TPP1-S^90-334^ or TPP1-L^1-334^. 10 pM ^32^P-end labeled GGTTAGGGTTAG DNA oligonucleotide was used in all binding experiments. See also Figure S2.

### Both TPP1-S and TPP1-L protect chromosome ends

TPP1 is unique among telomeric proteins as it not only recruits telomerase to telomeres for end replication, but it also helps protect chromosome ends (Palm and de Lange, 2008). To examine if the short and long isoforms of TPP1 differ in their end protection functions, we first asked how TPP1-S and TPP1-L M87A interact with their shelterin binding partners POT1 and TIN2. Co-immunoprecipitation experiments showed that both FLAG-tagged TPP1-S and TPP1-L M87A efficiently pulled down transiently co-expressed POT1 or TIN2 on anti-FLAG conjugated beads (Fig. S2C,D). Additionally, both the TPP1-S/TIN2 and the TPP1-L M87A/TIN2 complexes bound FLAG-POT1 to similar extents, ruling out any major differences in how the two TPP1 isoforms integrate into the shelterin complex (Fig. 2C). Furthermore both purified TPP1^90-334^ (TPP1-S lacking C-terminal domain) and TPP1-L^1-334^ (TPP1-L lacking C-terminal domain) protein enhanced POT1’s ability to bind telomeric DNA in a quantitative filter-binding assay (Fig. 2D,E). In agreement with these data, both TPP1-S and TPP1-L constructs co-localized with telomeric DNA (Fig. S2E) and neither isoform gave rise to substantial number of telomere-dysfunction induced foci when overexpressed (>70% of cells had no TIFs) (Fig. S2F). Taken together these data strongly suggest that both TPP1-L and TPP1-S are fully proficient at end protection.

### TPP1-L blocks telomere extension by telomerase

Given that both TPP1 isoforms are indistinguishable in their ability to protect chromosome ends or recruit telomerase to telomeres, we asked if the lack of telomere elongation in TPP1-L M87A overexpressing cells was indicative of a telomerase processivity defect. For this, we performed direct telomerase primer extension assays using purified POT1, TPP1^90-334^, and TPP1^1-334^ proteins with super telomerase extracts from cultured HEK 293T cells (Cristofari and Lingner, 2006). We measured telomerase processivity over time and found that at early time points TPP1^1-334^ was modestly, yet consistently, defective in stimulating telomerase processivity compared to TPP1^90-334^ (Fig. 3A; two additional replicates in Fig. S3A,B). However under the primer extension assay conditions, the TPP1^1-334^ protein underwent proteolytic trimming at the N-terminus (Fig. 3B), which could relieve telomerase inhibition and lead to an underestimation of the telomerase processivity defect (Fig. 3B).

**Figure 3.**
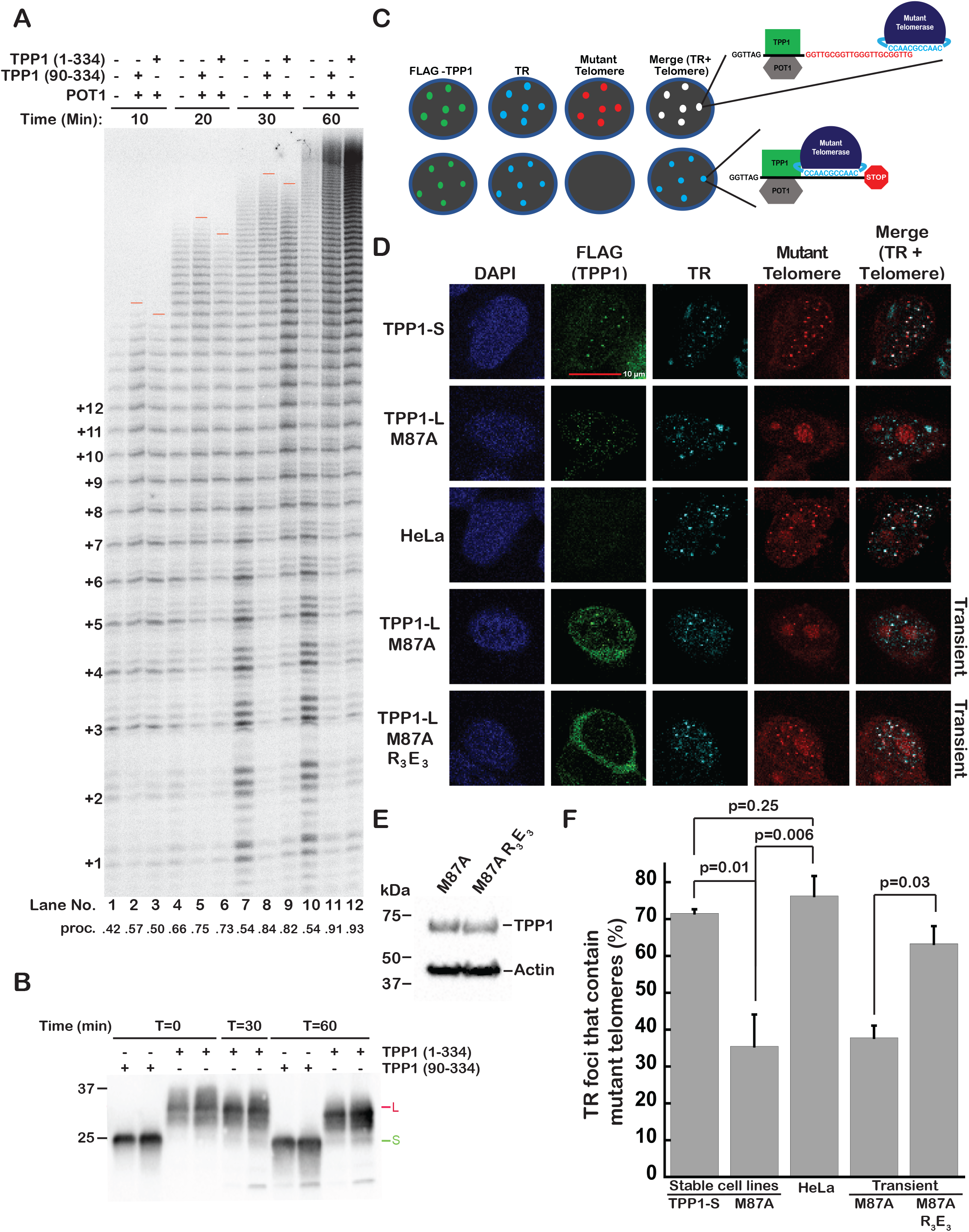
TPP1-L is deficient in activating telomere synthesis following telomerase recruitment. (A) Direct telomerase activity assay with purified POT1, TPP1-S^90-334^, and TPP1-L^1-334^ proteins at 30°C for the indicated incubation times. Red horizontal bars approximately mark the longest detectable product. The ratio of the total intensity from bands representing the addition of 9 or more hexad repeats over the total intensity of the lane is shown below as processivity (proc.). (B) Anti-TPP1 immunoblot of telomerase activity reactions (minus radiolabeled dGTP) at the indicated time-points. Green and red bars indicate the center of the TPP1-S and TPP1-L proteins, respectively, at time = 0. (C) Cartoon representation of expected outcomes of the *in vivo* IF/co-FISH telomere extension assay involving “FLAG-TPP1” (green), “TR” (cyan), and newly synthesized mutant telomeric repeats “Mutant Telomere” (red). White foci in “Merge” of TR and mutant telomere signals depicts colocalization. (D) HeLa-EM2-11ht cells transiently transfected with WT TERT and mutant telomerase RNA (TR) that contains a non-telomeric GCCAAC (WT: CCAAUC) template sequence. Immunofluorescence/co-fluorescence *in situ* hybridization was used to visualize telomere extension by telomerase in parental HeLa-EM2-11ht cells, indicated stable cell lines, and transiently transfected cells. “FLAG-TPP1” and “TR” were visualized as in Figure 2. “Mutant telomere” refers to signal from a Cy3-labled PNA probe against the mutant telomere sequence (red) that was synthesized by telomerase containing a mutant RNA template. Non-specific staining of the nucleolus by the mutant telomere probe occurred in all cells including HeLa cells that were not transfected with mutant telomerase (data not shown). (E) Immunoblot of lysates from HeLa-EM2-11ht cells transiently transfected with FLAG-TPP1-L M87A or FLAG-TPP1-L M87A-R_3_E_3_. (F) Quantitation of *in vivo* telomere extension data for which (D) is representative. For each experiment >300 TR foci were scored and the mean percent (bar) and standard deviation (error bar) of TR foci that contain mutant telomeres were plotted for triplicate measurements. Statistical significance was scored using a two-tailed student’s t test. See also Figure S3.

To circumvent the limitations posed by TPP1^1-334^ protein expressed recombinantly in *E. coli*, we utilized an in-cell telomere extension assay to visualize newly synthesized telomeric repeats (Fig. 3C) (Diolaiti et al., 2013). We transiently transfected HeLa-EM2-11ht cells and stable cell lines with WT TERT and a mutant telomerase RNA (TR) that contains a non-telomeric GCCAAC (WT: CCAAUC) template sequence as previously described. This system allows us to simultaneously determine the localization of FLAG-TPP1 protein (green; Fig. 3C) by IF, as well as mutant telomerase (telomerase^mutTR^; cyan; Fig. 3C) and newly added mutant telomere repeats (red; Fig. 3C) by FISH. We envisioned two contrasting scenarios. First, the TPP1 isoform will recruit telomerase^mutTR^ to the telomere where it will synthesize new mutant repeats (Top; white spots for TR + mutant telomere; Fig. 3C). Second, the TPP1 isoform will recruit telomerase^mutTR^ to the telomere, but telomerase^mutTR^ will be unable to synthesize mutant repeats (Bottom; cyan spots for TR + mutant telomere; Fig. 3C). As expected, TPP1-S overexpression resulted in telomerase^mutTR^ recruitment to telomeres and significant incorporation of newly synthesized mutant repeats, with more than 70% of TR foci containing mutant telomeres (Fig. 3D,F). In contrast, overexpression of TPP1-L M87A resulted in significantly fewer foci with mutant repeats, with only 35% of TR containing mutant telomere signal above background (Fig. 3D,F). The mutant telomere foci that were visible in TPP1-L M87A cells were also noticeably dimmer than those in TPP1-S overexpressing cells. As in TPP1-S overexpression cells, over 75% of telomerase^mutTR^ in HeLa cells (lacking TPP1 overexpression) contained mutant telomeres (Fig. 3D,F). Therefore the striking TPP1-L phenotype cannot be attributed to differences in TPP1-L and TPP1-S overexpression levels, and instead suggests that TPP1-L blocks telomerase action at telomeres.

To examine how TPP1-L restricts telomerase action, we looked more closely at the 86 amino acids unique to TPP1-L. This region is extremely basic (theoretical pI: 12.3) and predicted to be unstructured, consistent with its glycine/proline/arginine-rich composition (Fig. 1B), our circular dichroism results (Fig. S3D), and the propensity of TPP1^1-334^ to undergo proteolysis (Fig. 3B) and aggregation (data not shown) *in vitro*. To test whether basic residues in the N-terminus of TPP1-L contribute to its ability to block telomerase activity, we mutated three arginine residues in TPP1-L to glutamate residues (R43E/R46E/R48E; TPP1-L M87A R_3_E_3_; red “R” residues in Fig. 1B; Fig. 3E). Transient transfection of TPP1-L M87A R_3_E_3_ rescued telomerase activity with 63% of telomerase^mutTR^ foci containing mutant telomere sequence, compared to only 37% with transient transfection of TPP1-L M87A (Fig. 3D,F). This result raises the possibility that TPP1-L aa 1-86 interacts with nucleic acid, however we did not observe any binding of TPP1-L to telomeric single-stranded DNA (with or without POT1; Fig. 2E) or differences in TERRA (noncoding RNA of telomeric sequence transcribed from subtelomeric promoters (Azzalin et al., 2007)) accumulation at TPP1-L versus TPP1-S telomeres (Fig. S3C). Regardless, our rescue does verify the separation-of-function between human TPP1 isoforms such that only TPP1-S can activate telomere synthesis.

### TPP1-L and TPP1-S transcripts are both primed but only TPP1-S accumulates in most cells

The GENCODE transcript set describes two distinct transcripts for the human *ACD* locus that differ only at their 5’ ends (Fig. 4A) (Harrow et al., 2012). The shorter transcript (referred to here as TPP1-S mRNA) initiates downstream of the codon for TPP1 Met1 and can be translated into TPP1-S protein (starting at Met87) but not TPP1-L protein. The longer transcript (referred to here as TPP1-L mRNA) can code for both TPP1-L and TPP1-S proteins. Genome-wide ChIP studies in 91 cell lines, including cancer cell lines, fibroblasts, and embryonic stem cells, show a bimodal enrichment of transcription factors at the 5’ ends of TPP1-L and TPP1-S transcripts indicative of two distinct promoters (Fig. 4A; Fig. S4A) (Gerstein et al., 2012; Wang et al., 2012; Wang et al., 2013). A similar accumulation of active chromatin marks is observed at both transcription start sites (Fig. S4A). GRO-cap and GRO-seq data, which capture nascent RNA, similarly show that both TPP1-L and TPP1-S promoters fire in the cell lines tested (Fig. 4A; Fig. S4A) (Core et al., 2014; Core et al., 2008). In contrast to these data, RNA-seq data suggest that the steady state level of TPP1-S mRNA is significantly higher than that of TPP1-L mRNA (10-100 fold more reads common to both isoforms than reads specific for TPP1-L; Fig. 4A; Fig. S4A) (Consortium, 2012) (data deposited by Wold Lab at Caltech). In fact, very few TPP1-L specific reads are visible above background. CAGE data, which capture the 5’ ends of capped RNA in cells (Carninci et al., 1996), reveal only a small number of reads for the 5’ ends of TPP1-L relative to TPP1-S mRNA (note that y-axis is in log scale; Fig. 4A), suggesting that although both transcripts are primed for transcription, only the short (TPP1-S) transcript accumulates at steady state. The underrepresentation of TPP1-L by RNA-seq and CAGE was observed in all the cell lines tested, including several cancer cell lines, and embryonic stem cells (Fig. 4A; Fig. S4A). The high ratio of TPP1-S:TPP1-L mRNA correlates with a high ratio for their rate of protein translation, as ribosome profiling reads for the two isoforms mirror the RNA-seq data in all of the cell lines tested (Fig. 4A; Fig. S4A) (Michel et al., 2014).

**Figure 4.**
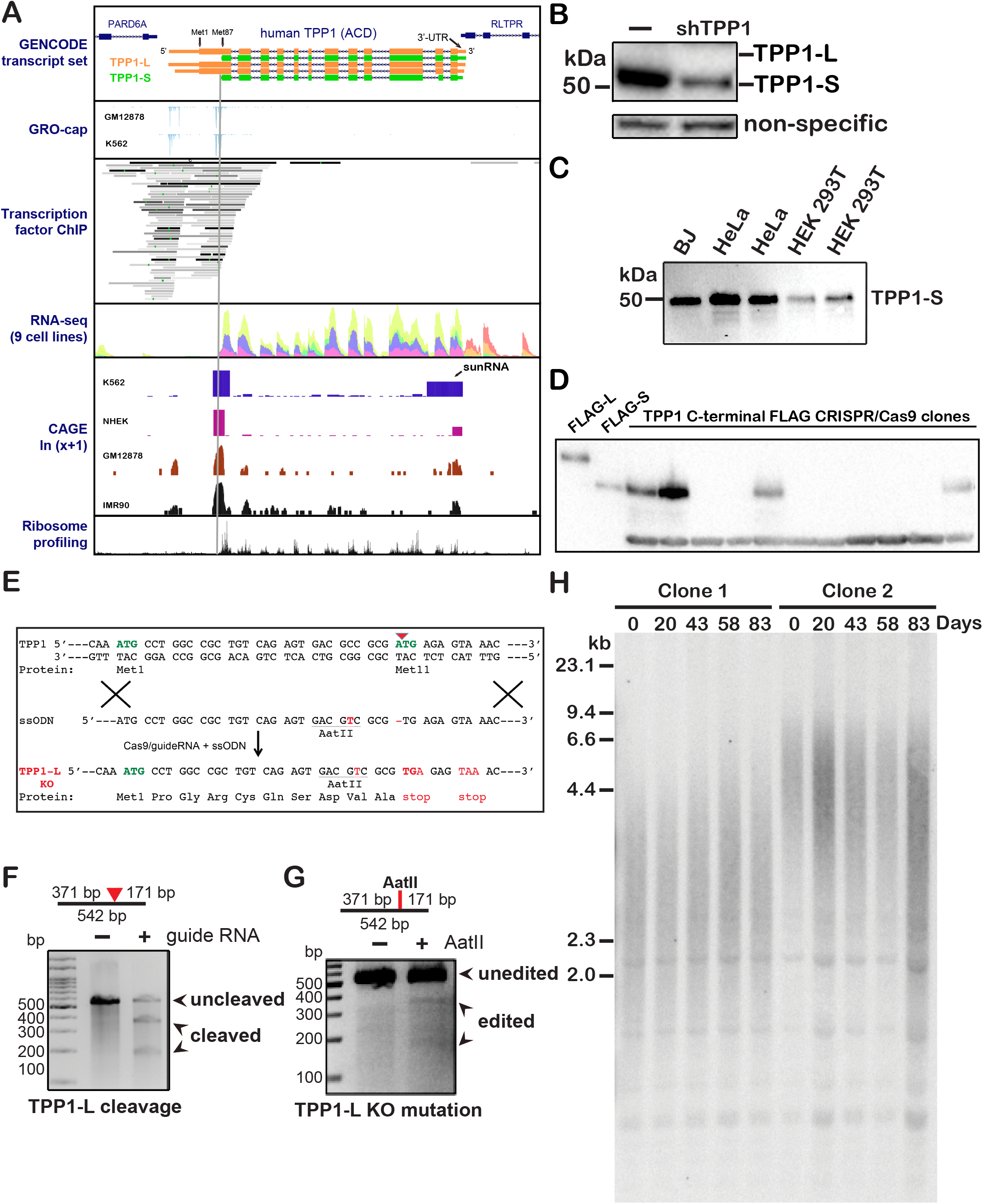
Separate promoters initiate TPP1-L and TPP1-S expression, but only TPP1-S accumulates in human cell lines. (A) Existing high-throughput sequencing information for the *ACD* locus was visualized using the UCSC genome browser. GENCODE reveals two major variations of transcripts differing only in their 5’ definition. We name these two mRNA isoforms TPP1-L mRNA and TPP1-S mRNA. The vertical grey bar denotes the 5’ end of TPP1-S mRNA. GRO-cap data for indicated cell lines show similar nascent transcription activity for TPP1-L and TPP1-S mRNA. Cumulative transcription factor ChIP data from a large number of human cell lines reveals two distinct peaks, suggesting independent transcription of TPP1-L and TPP1-S mRNA from two dedicated promoters. Cumulative RNA seq from nine cell lines (GM12878, H1-human embryonic stem cell line, HeLa-S3, HepG2, HSMM, HUVEC, K562, NHEK, and NHLF) shows very few reads specific to TPP1-L compared to TPP1-S mRNA at steady-state. Note that the cumulative plot is fully representative of data from each individual cell line (see Fig. S4A). CAGE data plotted in natural logarithm scale capturing capped RNA 5’ ends is shown for the polyA enriched nuclear fraction of the indicated cell lines. CAGE tags at the 3’ end of TPP1 mRNA indicative of the 5’ end of the sunRNA are indicated with an arrow. Cumulative ribosome profiling (ribo-seq) data obtained for a large number of human cell lines spanning twelve independent studies is shown. The cumulative plot is fully representative of data from each individual cell line (see Fig. S4A for data from an embryonic stem cell line). (B) Validation of rabbit polyclonal antibody against human TPP1 protein (epitope is in the OB domain; Bethly Labs) using a previously characterized shRNA against TPP1 in HeLa-EM2-11ht cells confirms presence of TPP1-S but not TPP1-L based on the expected size (marked on right). (C) Pull down of endogenous TPP1 from indicated cell types. A biotinylated telomeric DNA oligonucleotide bound to POT1 and immobilized on streptavidin beads was incubated with lysates from HeLa, HEK 293T, or BJ cells to enrich for TPP1. After wash steps, the beads were subjected to an immunoblot using a rabbit anti-TPP1 antibody (Bethyl lab). (D) Lysates from clonal HEK 293T cell lines with endogenous TPP1 tagged C-terminally with 3X-FLAG sequence using CRISPR-Cas9 technology were subjected to anti-FLAG immunoblotting. “FLAG-L” and “FLAG-S” indicate control lysates of HeLa-EM2-11ht cells transiently transfected with plasmids encoding FLAG-TPP1-L and FLAG-TPP1-S, respectively. (E) Strategy used to engineer TPP1-L KO cells. A guide RNA was designed to cleave the ATG codon for M11(at the position indicated by the red arrowhead), the only methionine between M1 and M87. A repair ssODN was designed to delete the “A” in the M11 ATG codon, creating an in-frame stop codon (TGA), and introducing an *Aat*II site using silent mutations. (F) A PCR-based Surveyor assay was performed to assess the efficiency of cleavage at the M11 codon by Cas9. The bar at the top shows the predicted product sizes upon Cas9-mediated cleavage, and the arrowheads alongside the gel indicates the un-cleaved and cleaved PCR products. (G) An *Aat*II digest performed two days post-transfection indicated a detectible level of edited cells in the population. (H) TRF Southern blot analysis of genomic DNA from CRISPR-Cas9 derived HeLa-EM2-11ht clones successfully knocked out for TPP1-L (with intact TPP1-S) for the indicated number of days in culture. See also Figure S4.

We independently explored TPP1 isoform abundance using immunoblotting with a well-characterized rabbit antibody that we have previously validated for detecting endogenous TPP1 (Bethyl Labs; A303-069A) (Bisht et al., 2016). The antibody detects a discrete band for TPP1-S (close to 50 kDa) that is diminished in the presence of an shRNA that targets a region common to both isoforms (Fig. 4B) (Bisht et al., 2016). We were only able to detect the TPP1-S isoform (Fig. 4B) even when we enriched for endogenous TPP1 in BJ fibroblasts, HeLa, and HEK 293T cell lysates using POT1-biotinylated DNA immobilized on streptavidin beads (Fig. 4C). To circumvent our inability to detect TPP1-L because of the sensitivity limit of the TPP1 antibody, we engineered a 3X-FLAG tag at the C-terminus of the endogenous *ACD* locus in HEK 293T cells using CRISPR-Cas9 technology (Fig. S4B) and isolated several clones that were accurately edited. In whole cell lysates (Fig. 4D) or after enrichment on anti-FLAG M2 affinity gel resin (Fig. S4C), we observed a band for TPP1-S but not TPP1-L in all the clones we tested. To further determine what effect potentially undetectable amounts of TPP1-L protein might have on telomere length maintenance, we used CRISPR-Cas9 technology to engineer TPP1-L knockout (KO) clonal HeLa cell lines. We targeted the endogenous *ACD* locus to generate a stop codon in the place of Met11 (the only Met between Met1 and Met87), preventing translation of TPP1 polypeptide from a Met upstream of aa 86, while preserving Met87 (Fig. 4E). We observed both efficient cleavage of the *ACD* locus (Fig. 4F) and proper editing thereafter (restriction cleavage at engineered *Aat*II site; Fig. 4G). We successfully isolated and propagated TPP1-L KO clones (Fig. S4D) but failed to observe any robust trend in telomere length, suggesting that HeLa cell telomere length is unaffected by KO of TPP1-L (Fig. 4H). In summary, although both TPP1-L and TPP1-S transcripts are similarly primed for transcription, TPP1-S is the major functional isoform in all the cell lines that were examined.

### An intragenic noncoding RNA in the 3’-UTR of *ACD* shuts down TPP1-L but not TPP1-S

Given that both TPP1-L and TPP1-S promoters displayed comparable potential for mRNA production, we wished to determine how TPP1-L mRNA is selectively down regulated in most human cells. We noted that the short (93 nt) 3’-UTR of the gene (*ACD*) coding for human TPP1 contains a stretch of 26 nucleotides that are almost fully complementary to regions in the TPP1-L open reading frame (ORF) (Fig 5A). Furthermore, the presence of CAGE tags close to the 3’ end of the *ACD/TPP1* gene were suggestive of ncRNA that encompasses the 3’-UTR sequence of TPP1 (see CAGE sub-panel in Fig. 4A). To verify that such a 3’-UTR derived ncRNA exists and examine what influence it might bear on TPP1-L expression, we performed 5’ RACE in HEK 293T cells. We identified a nested set of ncRNAs with the complete 3’-UTR sequence but differing at their 5’ ends, in agreement with the broad peaks in the CAGE data (Fig. 5B; Fig. 4A). We call these ncRNAs <u>s</u>ilencing 3’-<u>U</u>TR-derived <u>n</u>oncoding RNAs or sunRNAs, based on their function described later.

**Figure 5.**
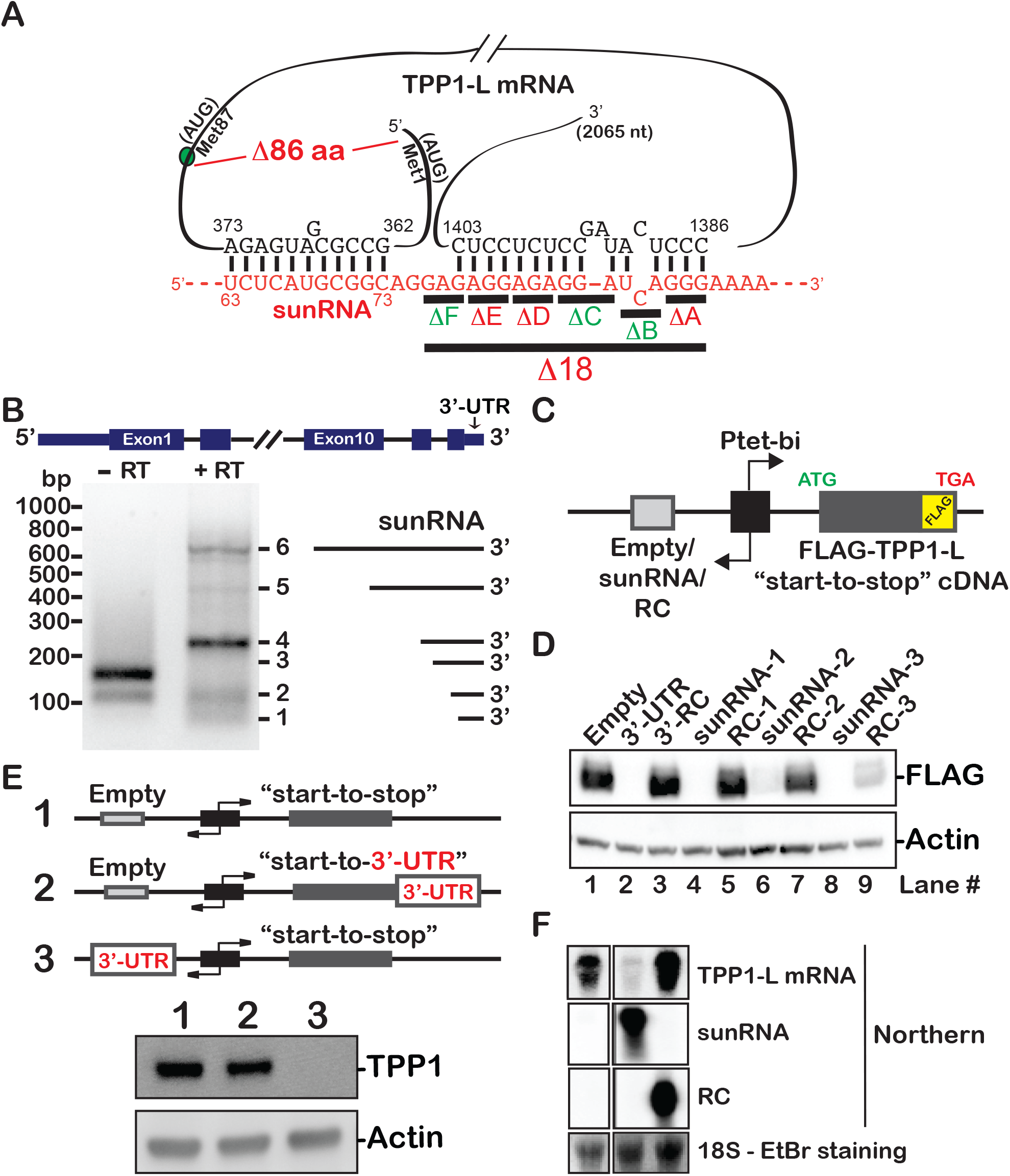
An intragenic RNA in the 3’-UTR of TPP1 completely shuts down TPP1-L. (A) Extensive complementarity was detected between two regions in the TPP1-L ORF and 3’-UTR (sunRNA). TPP1-S mRNA lacks the sequence that is complementary to the 63-73 nt region of 3’-UTR. Three nucleotide deletions were engineered in the sunRNA. Deletions in red are in regions of perfect predicted complementarity between the sunRNA and TPP1 mRNA, while deletions in green are in regions where the predicted complementarity is imperfect. (B) TPP1 sunRNAs were cloned from HEK 293T cells using 5’ RACE. The heterogeneity in length and 5’ end definition, as depicted to the right of the gel, is consistent with the broad nature of the sunRNA peaks in the CAGE data. (C) Schematic of the vector used to study the effect of sunRNA expression on TPP1 protein. Expression of both the protein and RNA constructs is driven simultaneously and from the same bi-directional promoter using doxycycline (“Tet-on” system). “RC” is a reverse complement of the sunRNA. (D) Western blot showing the silencing of TPP1-L by all the sunRNA constructs that were tested. “3’-UTR” indicates a minimal sunRNA-like construct that contains only the 93 bp between the stop codon and the start of the polyA tail of TPP1 mRNA. (E) The 3’-UTR sequence of TPP1 silences TPP1-L when expressed as a ncRNA (in *trans*), but not when it is inherent to TPP1-L mRNA (in *cis*). (F) Northern blot analysis of TPP1-L mRNA, sunRNA, and RC in the indicated transfections. 18S signal from ethidium bromide staining was used as a loading control. See also Figure S5.

We studied how the sunRNA affects TPP1-L levels using a bidirectional expression plasmid to co-express a protein-coding cDNA and a ncRNA gene as described previously (Bisht et al., 2017; Nandakumar et al., 2012). We cloned FLAG-TPP1-L cDNA (“start codon to stop codon” cDNA) in one multiple cloning site, and the cloned intragenic sunRNA candidates in the other (Fig. 5C). A plasmid encoding FLAG-TPP1-L, but lacking the sunRNA (Empty) was engineered as a negative control. Strikingly, while expression of FLAG-TPP1-L/Empty resulted in the expected accumulation of FLAG-TPP1-L protein (lane 1; Fig. 5D), co-expression of FLAG-TPP1-L and the sunRNA completely abrogated detection of FLAG-TPP1-L protein (Fig. 5D; lanes 4, 6, & 8). sunRNA-1, sunRNA-2, and sunRNA-3 were all able to shutdown TPP1-L expression, suggesting that the 3’-UTR sequence common to them is important for this process. Indeed, the 93 nt TPP1 3’-UTR RNA common to all sunRNAs was sufficient to silence TPP1-L (Fig. 5D; lane 2). To demonstrate that silencing of TPP1-L is specific to the sunRNA we designed plasmids encoding FLAG-TPP1-L and the reverse complement sequence (RC) of each sunRNA, so as to conserve the RNA length, GC content, and (to a large extent) secondary structure, but not sequence. These control constructs resulted in an accumulation of TPP1-L protein similar to that of FLAG-TPP1-L/Empty (Fig. 5D; lanes 5, 7, & 9), suggesting TPP1-L silencing by the sunRNA is sequence specific. CAGE tags at the 3’ end of the *ACD* gene point to the sunRNA transcript being separate from the mRNAs encoding the TPP1 isoforms. Accordingly, TPP1-L was silenced when its 3’UTR was co-expressed in *trans* (#3 in Fig. 5E) but not when the 3’UTR was part of the TPP1-L mRNA (#2; Fig. 5E). Taken together, these data suggest that TPP1-L is silenced by a ncRNA that is derived from its own 3’-UTR.

We verified that TPP1-L protein expression levels correlated with mRNA abundance by Northern blot analysis, which depicted a clear loss of TPP1-L mRNA in the presence of the sunRNA, but not RC (Fig. 5F). Robust silencing was also recapitulated when TPP1-L and the sunRNA were expressed from separate plasmids (Fig. S5A) and when TPP1-L expression was regulated by a constitutively active CMV promoter (Fig. S5B). Finally, the silencing is TPP1 mRNA-specific, as the 3’-UTR ncRNA of TPP1 is unable to silence other shelterin components such as TIN2 or POT1 (Fig. S5C).

Intriguingly, the sunRNA contains a stretch of 10 nt that are fully complementary to a region at the 5’ end of the TPP1-L mRNA, but absent in the TPP1-S mRNA (Fig. 5A). To test the role of this selective complementarity in the silencing of TPP1-L versus TPP1-S, we co-expressed TPP1-S with the sunRNA. In sharp contrast to the complete silencing of TPP1-L, the sunRNA failed to diminish the levels of TPP1-S mRNA or protein (Fig. 6A,B). Because mouse TPP1 begins at a Met equivalent to human Met87 (Fig. 1B) it lacks the first target site for the sunRNA (red in Fig. S6A). However mouse TPP1 also lacks 29 aa that are present in both human TPP1-S and TPP1-L (aa 334-362) but which lie outside TPP1’s TERT, POT1, and TIN2 binding regions. Most remarkably, the mRNA that codes for these residues harbors a putative second target site of complementarity for the sunRNA (TPP1-L mRNA nt 1386-1403 shown in black in Fig. 5A and red in Fig. 6C and Fig. S6A). To test the importance of this potential target site, we created a series of deletion mutations within the sunRNA. Notably, only sunRNA constructs with deletions that significantly disrupted complementarity (i.e. 3 bp) lost silencing capacity and rescued TPP1-L expression (deletions marked in red in Fig. 5A and Fig. 6D). In contrast, sunRNA constructs with deletions disrupting less than 3 bp of complementarity continued to silence TPP1-L expression (deletions marked in green in Fig. 5A; Fig. 6D). Furthermore, neither the human TPP1 sunRNA nor the 3’-UTR of mouse TPP1 is able to knockdown expression of mouse TPP1, further confirming that this phenomenon is absent in mice (bottom image of Fig. 6E: compare lanes 2 and 3; compare lanes 5 and 6).

**Figure 6.**
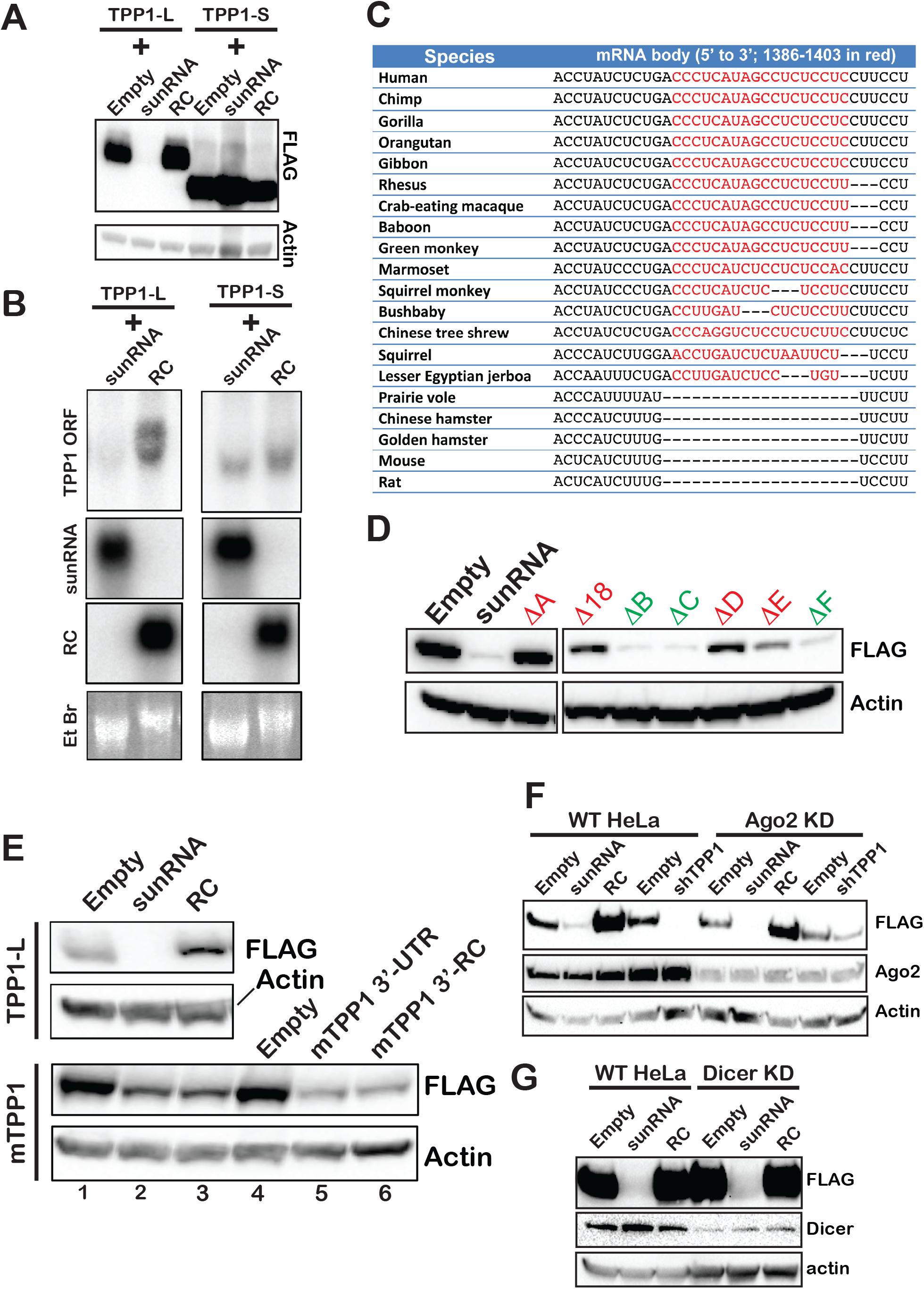
The sunRNA silences TPP1-L but not TPP1-S. (A) Parallel comparison of lysates from cells co-expressing the sunRNA with FLAG-TPP1-L or FLAG-TPP1-S reveals that the sunRNA does not silence TPP1-S. (B) Northern blot analysis confirms that the sunRNA diminishes TPP1-L mRNA levels but not TPP1-S mRNA levels compared to the RC control. (C) Sequence alignment of TPP1 homologs reveals a second primate specific sequence element in the TPP1 ORF (first element being aa 1-86 of TPP1-L) that is absent in rodent TPP1 proteins. This region includes the target site of the sunRNA (shown in red), but codes for amino acids that are not required for any of TPP1’s known telomeric functions. (D) Anti-FLAG immunoblot analysis of lysates from cells transiently transfected with sunRNA mutants (defined in Fig. 5A) show that disruption of the predicted complementarity between TPP1-L and the sunRNA reduces TPP1-L silencing. (E) *Top:* Replicate data for the sunRNA silencing FLAG-TPP1-L. *Bottom:* Bi-directional constructs for co-expression of FLAG-tagged mouse TPP1 with Empty vector (lane 1), human TPP1 3’-UTR (sunRNA; lane 2), human TPP1 RC (lane 3), Empty vector (lane 4), mouse TPP1 3’-UTR (lane 5), or the reverse complement (RC) of the mouse TPP1 3’-UTR (lane 6) were transfected into HeLa cells and analyzed using anti-FLAG immunoblotting. (F and G) Immunoblots from HeLa transfected with FLAG-TPP1-L and Empty/sunRNA/RC constructs; and siRNAs for Ago2 (F) or Dicer (G) knockdown. shTPP1 is the same construct used in Fig. 4B. Note that it is very effective in knocking down TPP1 compared to the Empty vector control in “HeLa”, but not in cells also knocked down for Ago2. See also Figure S6.

To further characterize the pathway through which the sunRNA inhibits TPP1-L mRNA we knocked down key proteins in the RNA interference pathway. Downregulation of *AGO2* and *DICER1* did not disrupt the silencing function, suggesting that the TPP1 sunRNA is not a cryptic microRNA (Fig. 6F,G). Similarly, knockdown of essential components of the nuclear or cytosolic RNA exosomes (involved in 3’ RNA degradation), decapping enzyme DCP2 (involved in 5’ mRNA degradation), XRN1 or XRN2 ribonucleases, or components of Lsm complexes (involved in splicing and RNA turnover) were unable to perturb TPP1-L silencing by the sunRNA (Fig. S6B-G). Together our data suggest a novel mechanism to selectively silence a specific protein-coding mRNA isoform using an intragenic ncRNA that we accordingly named <u>s</u>ilencing 3’-<u>U</u>TR-derived <u>n</u>oncoding RNA or sunRNA.

### TPP1-L expression switches on during spermatogenesis

TPP1-L is undoubtedly the minor TPP1 isoform in all the somatic cell lines, embryonic stem cell lines, and cancer cell lines that we analyzed. Yet the presence of an actively firing promoter for TPP1-L, the unique ability for TPP1-L to recruit but not activate telomerase, and the intricate sunRNA regulatory mechanism to shut down TPP1-L expression suggest that it is unlikely to be a vestigial protein. Because we were unable to detect TPP1-L in human cell lines, we shifted our focus to expression profiles in human tissues and organs. Inspection of previously archived RNA-seq data revealed that TPP1-S expression dominated in many human tissues/organs, including brain, lymph node, adipose tissue, breast, colon, and skeletal muscle, with very few TPP1-L-specific reads detected (Fig. 7A) (Burge lab data visualized on UCSC genome browser; (Wang et al., 2008)). However several reads specific to the first 86 aa of TPP1-L were detectible in testes, on par with some exons that are fully shared between TPP1-L and TPP1-S. To further verify this observation and to estimate the TPP1-L:TPP1-S ratio, we examined the Genotype-Tissue Expression (GTEx) database that quantitates isoform expression from exonic data in various human tissues (The Genotype-Tissue Expression (GTEx) Project was supported by the Common Fund of the Office of the Director of the National Institutes of Health, and by NCI, NHGRI, NHLBI, NIDA, NIMH, and NINDS. The data used for the analyses described in this manuscript were obtained from: the GTEx Portal on 05/10/18). The GTEx data for TPP1 expression in testes confirmed that TPP1-L is indeed the major isoform, expressing at ~10-fold over TPP1-S (Fig. 7B). As TPP1-L mRNA expression does not confirm TPP1-L protein production or preclude TPP1-S translation from the downstream Met87 start codon, we lysed normal human testes tissue and performed immunoblotting using a rabbit TPP1 antibody. Lysates from HeLa cells transfected with FLAG-TPP1-S and FLAG-TPP1-L served as markers on the blot. In sharp contrast to ribosome profiling data (Fig. 4A; Fig. S4A) and Western blotting analysis of somatic cells (Fig. 4B-D), the only detectible band in the chromatin enriched fraction of testes lysate corresponded to the length of TPP1-L (Fig. 7C). These data are fully consistent with TPP1-L accumulating specifically in testes.

**Figure 7.**
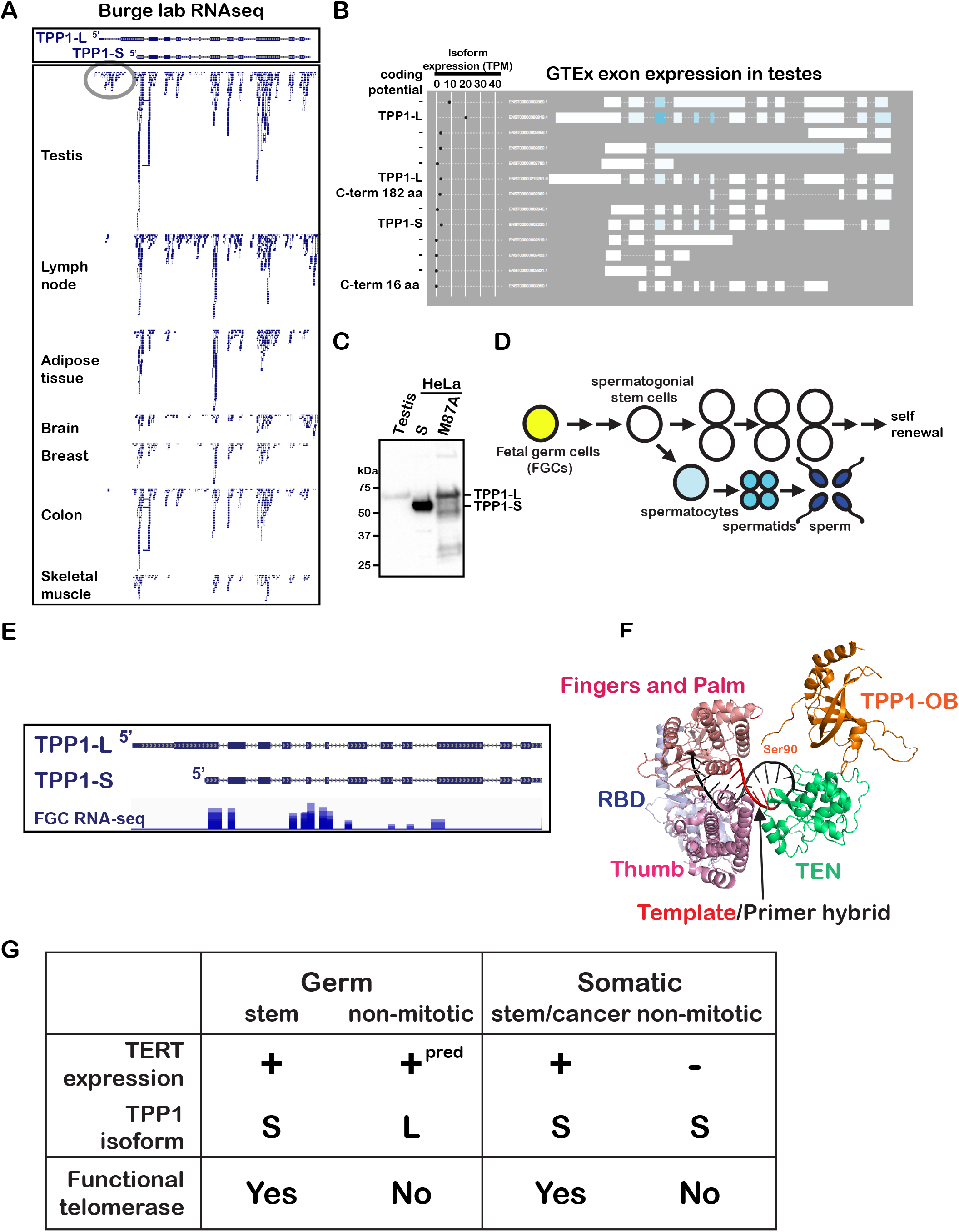
TPP1-L is robustly upregulated during spermatogenesis. (A) Deposited Burge lab RNA seq data from the indicated tissues/organs was visualized on the UCSC genome browser. The conspicuous abundance of TPP1-L specific reads in testes relative to other tissues is highlighted with a grey oval. (B) Tissue/Organ-specific isoform annotation by GTex transcriptome analysis reveals a greater abundance of TPP1-L than TPP1-S mRNA in testes. (C) Immunoblot analysis using the anti-rabbit TPP1 polyclonal antibody (Bethyl labs) of the chromatin-associated fraction of cells from a biopsy of a human testis reveals a band corresponding to TPP1-L but not TPP1-S. FLAG-tagged TPP1-L and TPP1-S overexpression cell extracts were run in parallel as size markers. (D) Schematic for the different stages of spermatogenesis. (E) Cumulative representation of published RNA-seq reads from 2,167 fetal germ cells (FGCs) aligned to the human *ACD* locus. (F) Structural model for telomerase recruitment to telomeres based on cryoEM structure of *Tetrahymena thermophila* telomerase (PDB accession: 6D6V) and *Tribolium Castaneum* TERT bound to a DNA-RNA hybrid (PDB accession: 3KYL). (G) Model in tabular form for how telomerase is regulated in various cell types depending on the TERT expression and TPP1 isoform status. “pred” indicates predicted TERT expression in differentiated germ cells in preparation of telomerase action in embryogenesis.

TPP1-L expression in the testes is likely representative of the large majority of the cells (~80%) that have undergone differentiation and maturation (Fig. 4D) (Fayomi and Orwig, 2018). Although there is insufficient RNA-seq data for (the more rare) human spermatogonial stem cells to make direct conclusions about TPP1 isoform expression in these cells (Hammoud et al., 2014), single-cell RNA-seq data for FGCs (precursors of adult spermatogonial stem cells) demonstrate that they express TPP1-S but not TPP1-L (Li et al., 2017) (Fig. 7E). Together our analysis of next-generation sequencing data for testes and cells therein provide compelling evidence that TPP1-S is the predominant isoform in the self-renewing germline population (spermatogonial stem cells), while TPP1-L dominates upon differentiation and maturation.

## Discussion

Our curiosity for the identities and functions of human TPP1 isoforms led to the discovery of two isoforms distinguished by their ability to activate telomere synthesis by telomerase and their distinct representation in somatic cells versus differentiated cells of the germline. We uncovered a novel regulatory ncRNA, which we call sunRNA, that dictates TPP1 isoform expression. Finally, our observation of an isoform switch in TPP1 unveils a new function for TPP1 specific to spermatogenesis, prompting the investigation of alternative isoforms of other telomeric and non-telomeric proteins in this unique stage of reproductive biology.

Our telomerase recruitment analysis, along with previously published reports, demonstrates clearly that TPP1-L is proficient at recruiting telomerase to telomeres. This suggests that both the TEL patch and NOB in TPP1-L are able to fully engage their target sites on TERT. However, telomerase recruited to the telomere by TPP1-L cannot efficiently extend chromosome ends. We therefore characterize TPP1-L as a telomerase activation-incompatible isoform of TPP1 that is distinct from all previously characterized TEL patch and NOB mutants of TPP1-S. To better understand how TPP1-L blocks telomerase action, we built a structural model of the human TPP1 OB domain bound to telomerase, based on the cryo-EM structure of *Tetrahymena thermophila* telomerase bound to the p50 protein (presumed homolog of human TPP1 based on its structure and function) (PDB accession: 6D6V) (Jiang et al., 2018) and the template-primer bound *Tribolium castaseum* TERT structures (PDB accession: 3KYL) (Mitchell et al., 2010) (Fig. 7F). In our model, the N-terminus of TPP1-S (Ser90; Fig. 7F) is in close proximity to the highly basic TEN domain of TERT that engages the template-primer hybrid. Based on the ability of TPP1 M87A R_3_E_3_ to rescue telomerase activation, we envision at least two possibilities for how TPP1-L inhibits telomere elongation. First, it is possible that electrostatic repulsion between the highly basic aa 1-86 of TPP1-L and TEN domain of TERT prevents telomerase from adopting a conformation that is conducive to processive telomere elongation. Second, it is possible that TPP1-L aa 1-86 directly binds to the telomeric DNA, TR, or the template-DNA hybrid, impeding proper engagement of the chromosome end by telomerase. However we did not observe any differences in the ability of TPP1-S or TPP1-L to associate with POT1-DNA *in vitro*, or TERRA (Azzalin et al., 2007)) in cells, suggesting that the differences in isoform function do not arise from interactions with these nucleic acids.

The separation-of-function for TPP1-S and TPP1-L isoforms implies that they have unique roles in distinct biological contexts. Analysis of high-throughput gene expression data led to the recognition of two dedicated promoters and transcripts for TPP1-L and TPP1-S protein production in human cells. The transcripts are not splicing isoforms, as they contain identical exon-intron junctions and sequences, but are rather alternative start site mRNAs. This results in the production of two protein isoforms, originating from a set of in-frame alternative start sites. Despite GRO-cap, GRO-seq, and transcription factor ChiP data confirming transcriptional activity at both promoters, we observed that TPP1-S is the main isoform in all somatic cells and in embryonic stem cells. The presence of TPP1-S in stem cells is fully consistent with their need to support telomerase activity for continued self-renewal (Fig. 7G). After differentiation, cells use a robust mechanism to switch off telomerase expression (Aubert, 2014; Gunes and Rudolph, 2013), rendering downregulation of TPP1-S (or upregulation of TPP1-L) unnecessary. Thus it is not surprising that TPP1-S is the major isoform in somatic cells, both before and after differentiation (Fig. 7G).

Telomerase is strictly regulated, with continued expression in actively dividing cells and robust shut down after cell differentiation. However spermatogenesis poses a unique challenge for telomerase regulation. Spermatogonial stem cells self-renew and thus it is not surprising that they are telomerase-positive (Izadyar et al., 2011). However, spermatogonial stem cells differentiate into spermatocytes, which undergo meiosis (single rounds of meiosis I and II) to form haploid spermatids and ultimately gametes (Fig. 7D) (Marston and Amon, 2004). These haploid cells are unique non-dividing cells that act as precursors to embryogenesis. They must block excessive telomerase action, much in the way differentiated somatic cells need to shutdown telomerase expression. Yet (unlike differentiated somatic cells) they must also be able to swiftly re-activate telomerase during embryogenesis, suggesting that there must be a mechanism to re-activate telomerase in the embryo after it is mitigated in the gametes. In the absence of any (known) non-pathological/natural mechanism to re-activate telomerase expression after it is shutdown, we propose that differentiated (i.e., non-mitotic) human germ cells retain telomerase expression but keep its activity in check by also upregulating TPP1-L (Fig. 7G). Indeed telomerase activity has been observed in human spermatogonial stem cells, but not in human sperm (Ozturk, 2015; Wright et al., 1996). Finally, upon fertilization, TPP1-S expression would resume, ensuring end replication (and protection) during embryogenesis and throughout life thereafter.

The differences in telomerase regulation between mice and humans are well documented, most often being attributed to the difference in total number of cell divisions between the organisms. In contrast to humans, mice express telomerase in most cells, including non-dividing somatic cells, making a check on telomere elongation in differentiated germ cells unnecessary (Prowse and Greider, 1995). Thus it is not surprising that TPP1-L is absent in mice. The absence of TPP1-L in mice also obviates the need for regulation using a sunRNA-mediated gene silencing mechanism (discussed below).

We have found that TPP1-L mRNA is selectively diminished by a novel intragenic RNA. Given the absence of transcription factor hotspots, GRO-cap/GRO-seq reads, and active chromatin marks, the TPP1 sunRNA appears to not be made *de novo* but to instead be derived from processing of the 3’ ends of TPP1-L and/or TPP1-S transcripts (Fig. 4A). Such processing has been observed for thousands of human transcripts, although the full functional implications of this phenomenon remain uncovered (Malka et al., 2017). However the sunRNA was identified from CAGE analysis of the polyA-enriched nuclear fraction, suggesting that it somehow undergoes maturation following its genesis from TPP1 mRNA in the nucleus. Based on its origin from the 3’ end of the TPP1 gene, the sunRNA can be placed in two additional ncRNA categories: “termini-associated sRNA” or TASR (Affymetrix and Cold Spring Harbor Laboratory, 2009; Kapranov et al., 2007) and a special variant of 3’ UTR-associated RNA or uaRNA (Mercer et al., 2011) that is not a *de novo* transcript.

To our knowledge, this is the first report of differential silencing of protein isoforms by an intragenic ncRNA. We show that two essential sequence elements in the TPP1 sunRNA are complementarity to two separate target regions in the TPP1-L mRNA sequence. The upstream target region is present only in TPP1-L, and absent in TPP1-S, and appears to be the basis for the isoform specificity. The downstream target region, shared by both human TPP1-S and TPP1-L, codes for a stretch of seemingly unstructured amino acids with no known function. Both of these sunRNA-recognition regions are conserved among primates but absent in the mouse TPP1 mRNA, suggesting that they play an important regulatory function at the mRNA level. We propose a model whereby TPP1-L mRNA is primed in all human cells, but it is shut down by the TPP1 sunRNA in somatic cells, allowing TPP1-S to emerge as the major isoform (Fig. 7G). The switch that regulates sunRNA levels during spermatogenesis remains to be established, pending further experimentation including CAGE analysis in cells of the testes.

There are several conceivable advantages of sunRNA-like intragenic regulatory mechanisms in complex genomes. (1) The *cis* nature of the transcription start sites of the silencer RNA and the target mRNA could provide for an easier “search” for the complementary sequence due to enhanced local concentrations of the two RNAs. This is particularly relevant in complex mammalian transcriptomes where RNA-RNA recognition in *trans* is entropically disfavored without involvement of specialized machineries such as in RNAi. It is interesting to note that in less complex genomes such as those in bacteria, ncRNAs called sRNAs expressed from the 3’-UTR of protein-coding genes, can silence mRNA targets in *trans* using sequence-complementarity driven mechanisms (Kunne et al., 2014; Miyakoshi et al., 2015; Waters and Storz, 2009). It is therefore tempting to speculate that sRNA-mediated mechanisms were largely lost (and replaced by RNAi) when more complex transcriptomes evolved, and only cis-sRNA (like sunRNA) mediated mechanisms were conserved due to proximity between the silencer and the target. (2) sunRNAs can utilize the mRNA polyadenylation signal for their own 3’ end maturation thus obviating the need for further 3’ maturation. (3) The selection pressure to conserve essential mRNA sequences will also help preserve the corresponding sunRNA sequence. Therefore, we hypothesize that the overlapping nature of mRNA and sunRNA has prevented identification of other sunRNAs, and many such ncRNAs and their associated biological functions await discovery.

## Acknowledgements

J.N. would like to thank Dr. Thomas Cech for providing him (during his stay in Dr. Cech’s lab) the freedom to explore ideas outside that of the main project, one of which paved the path to sunRNA discovery. We would like to thank Dr. Peter Baumann for gifts of generous aliquots of several siRNAs and antibodies (detailed in STAR Methods and Resource Table); Dr. Thomas J. Giordano and Michelle Vinco of the Tissue Core of University of Michigan Rogel Cancer Center for providing testes tissue; Dr. Gregg Sobocinski for help with microscopy; Dr. Sneh Lata for preliminary bioinformatics analysis; Dr. Ivan Maillard and Dr. Catherine Keegan for useful discussions; Dr. Sue Hammoud for providing insights into spermatogenesis; and members of the Nandakumar laboratory for critical feedback. This work was funded by NIH Grants R01GM120094 (to J.N.), R01AG050509 (to J.N.; co-PI), and T32GM007544 (to University of Michigan Genetics Training Program; S.G. was a trainee); and the American Cancer Society Research Scholar grant 130882-RSG-17-037-01-DMC (to J.N.).

## Author contributions

J.N. conceived the hypothesis surrounding TPP1-L and TPP1-S separation-of-function and differential silencing by sunRNA. J.N., S.G., and K.B. designed experiments. S.G. performed all experiments involving TPP1-L (including TPP1-L M87A) and TPP1-S expression, separation-of-function, CRISPR KO, and localization. S.G. was assisted by V.M.T. in telomerase primer extension experiments and purification of POT1 and TPP1 proteins. K.B. performed all sunRNA-related experiments and knock-in of the C-terminal FLAG tag at the endogenous locus for TPP1 expression. C.J.S. performed bioinformatics analysis of data in the SRA. All authors analyzed the data, and S.G. and J.N. wrote the manuscript with detailed feedback from V.M.T.

## DECLARATION OF INTERESTS

The authors declare no competing interests.

## STAR Methods

### Molecular Cloning and site-directed mutagenesis

3X-FLAG-tagged constructs and 6x-MYC-tagged constructs for human cell expression were cloned into the pTET-IRES-eGFP-Bi4 vector; and 3X-FLAG-TPP1-L and 3X-FLAG-TPP1-L-M87A constructs were cloned into p3X-FLAG-TPP1-F3 vectors as previously described(Grill et al., 2018; Nandakumar et al., 2012). Additionally, 3X-FLAG-TPP1-S and 3X-FLAG-TPP1-L M87A constructs were cloned into a pcDNA3 derived vector and used to generate the M87A R_3_E_3_ construct for transient transfection in HeLa cells. All TPP1-S constructs used for human cell expression contained cDNA sequences corresponding to aa 87-544 of the TPP1 protein, while all TPP1-L constructs contained cDNA sequences corresponding to aa 1-544. The M87A mutation in the TPP1-L expression plasmid was introduced using the QuikChange® Site-Directed Mutagenesis Kit (Agilent Technologies) and complementary mutagenic primers (Integrated DNA Technologies). The resulting TPP1-M87A plasmid was sequenced to confirm both the presence of the intended mutation and the absence of unwanted errors introduced during PCR amplification/cloning. All additional site-directed mutagenesis, including mutation of TPP1-L M87A to M87A R_3_E_3_, and mutation of TR to mutant TR was performed using the same protocol as above. For bacterial expression of TPP1^90-334^ and TPP1^1-334^, the appropriate TPP1 sequence was cloned into the pET-Smt3 vector using standard restriction endonuclease cloning as described previously (Grill et al., 2018). For insect cell expression and purification of human POT1 the pFBHTb-Smt3star-hPOT1 was used as described previously (Kocak et al., 2014).

### Cell culture

Cell culture was performed as described previously (Bisht et al., 2016; Grill et al., 2018). HeLa-EM2-11ht cells were used in all transient transfection, while both HeLa-EM2-11ht and HEK 293T cells were used in CRISPR-Cas9 experiments. All cells were cultured at 37°C in the presence of 5% CO2 and propagated in growth medium containing modified DMEM (Gibco; Dulbecco’s Modified Eagle Medium; 11995-065), 100 U/mL penicillin, 100 µg/mL streptomycin, and 10% FBS. For all experiments requiring induction, doxycycline was added to a final concentration of 200 ng/mL to drive a tetracycline-inducible promoter within the p6X-MYC-BI4 or p6X-FLAG-BI4 plasmids (FLAG-TPP1-S, FLAG-TPP1-L, FLAG-TPP1-L M87A, ΔNOB).

### Generation of stable cell lines expressing TPP1-L and TPP1-L M87A

Stable cell line generation was performed exactly as described previously (Grill et al., 2018). Briefly, HeLa-EM2-11ht cells (Weidenfeld et al., 2009) were co-transfected with 1 µg each of the p3X-FLAG-TPP1-F3 (TPP1-L or TPP1-M87A) and Flp recombinase-expressing plasmid (that also codes for puromycin resistance). Cells were selected for one day using puromycin (5 µg/ml; Sigma-Aldrich), and then fresh medium with ganciclovir (50 µM; Sigma-Aldrich) was added for 10 days of negative selection. Individual clones were picked and expanded, and positive clones were selected based on Western blot analysis of FLAG-TPP1 signal after overnight induction with doxycycline (200 ng/ml). Generation of the stable cell lines expressing vector, TPP1-S, and TPP1ΔNOB constructs used in this study have been described previously (Grill et al., 2018; Nandakumar et al., 2012)

### Telomere restriction fragment length analysis

Telomere length analysis was performed as described previously (Grill et al., 2018; Nandakumar et al., 2012). Briefly, genomic DNA from stable cell lines expressing FLAG-TPP1 constructs (TPP1-S/TPP1-L/TPP1-M87A) and a vector control was isolated from confluent 6 cm dishes using the GenElute kit (Sigma). 2 µg of DNA was digested with *Hinf*1 and *Rsa*1 and incubated overnight at 37°C. The digested DNA was run on a 0.8% 25 cm long Agarose-1X TBE gel along with a Lambda DNA-*Hind*III digest ladder (NEB) at a constant 50 V for 20-23 h. After imaging with a florescent ruler, the gel was transferred to a sheet of dry Whatman filter paper and dried at 55°C for one h. After drying, the filter paper was removed and the gel was soaked in buffer containing 0.5 M NaOH for 30 min, rinsed with water, and shaken in a solution of 0.5 M Tris-Cl and 1.5 M NaCl (pH 7.5) for 30 min. The gel was then prehybridized in Church buffer (0.5 M sodium phosphate buffer [pH 7.2], 1% bovine serum albumin [BSA], 1 mM EDTA, 7% SDS) for 30 min at 65°C in a rotating hybridization oven. A telomeric probe of sequence (TTAGGG)_4_ was 5’ ^32^P-labeled with T4 PNK (NEB) and added at 20 million cpm to the gel. Hybridization continued overnight at 55°C. After overnight hybridization the gel was washed three times with 2X SSC for 10 min and exposed to a phophorimager screen for 24-72 h. The gel was analyzed using the Imagequant TL software and calibrated using the molecular weights of the lambda DNA ladder. The mean telomere length for each lane was plotted as a function of days in culture for each cell line. A linear regression (MS Excel) was used to calculate the rate of telomere elongation/shortening.

### Immunofluorescence-fluorescence *in situ* hybridization

#### TIF analysis (co-IF)

Co-IF experiments for TIF analysis were adapted from protocols described previously (Nandakumar et al., 2012). Briefly, co-IF was performed using stable cell lines expressing either FLAG-TPP1-S or FLAG-TPP1-L M87A. Cells were induced with doxycycline for 3 days and ~100,000 cells were seeded on coverslips in growth medium containing doxycycline. 24 hours after seeding, medium was removed, cells were washed with PBS, and all subsequent steps were performed at room temperature. Cells were fixed with 4% formaldehyde in PBS for 10 min and then washed in PBS before permeabilization with 0.5% Triton-X 100. Cells were blocked (PBS-T containing 3% FBS) for 30-60 minutes and incubated with mouse monoclonal anti-FLAG M2 antibody (Sigma; F1804; 1:500) and rabbit polyclonal anti-53BP1 antibody (Novus Biologicals; NB100-304; 1:1,000) diluted in blocking buffer. After 60 min of incubation, cells were washed with PBS and incubated with Alexa Fluor 568-conjugated anti-mouse IgG (Life Technologies) and Alexa Fluor 647-conjugated anti-rabbit IgG (Life Technologies) diluted 1:500 in blocking buffer for 30 minutes. Cells were mounted on microscope slides using ProLong Gold mounting medium with DAPI (Life Technologies). Coverslips were sealed with transparent nail polish and stored at 4°C until the time of imaging. All imaging was performed using a laser scanning confocal microscope (SP5; Leica, Germany) equipped with a 100X oil objective. ImageJ and Adobe Photoshop were used to process all images. Colocalizations of FLAG-TPP1 foci and 53BP1 foci were quantified manually by two separate individuals.

#### Telomerase recruitment to telomeres (IF-FISH)

IF-FISH experiments for telomerase recruitment were performed exactly as described previously (Grill et al., 2018). Briefly, IF was performed prior to FISH using mouse monoclonal anti-FLAG M2 (Sigma; F1804; 1:500) in combination with Alexa Fluor 568-conjugated anti-mouse IgG (Life Technologies) to visualize FLAG TPP1 proteins (FLAG-TPP1-S, FLAG-TPP1-L, FLAG-TPP1-M87A) by IF. A mixture of Cy5-conjugated probes complimentary to TR (Abreu et al., 2011) was used to detect TR by FISH. Imaging was performed as described above. Colocalizations were quantified manually by two separate individuals.

#### Telomere localization of TPP1-S and TPP1-L M87A (IF-FISH)

Telomere localization of all TPP1 constructs was performed exactly as previously described (NOB) (Grill et al., 2018). Briefly, a Cy3-conjugated PNA-(CCCTAA)_3_ probe was used to visualize telomeres by FISH. Cells were fixed, permeabilized, and soaked in 2X SSC, 50% formamide before being heated for 6 min at 80°C in the presence of the probe. After 2 h of hybridization cells were washed and subjected to immunofluorescence for FLAG-TPP1. Imaging was performed as described above.

#### TERRA localization at telomeres

This protocol was adapted from TERRA-FISH protocols previously published in the literature(Azzalin et al., 2007). 48 h after doxycycline induction, FLAG-TPP1-S and FLAG-TPP1-L M87A stable cell lines were washed with PBS and permeabilized in CSK buffer (100 mM NaCl, 300 mM sucrose, 3 mM MgCl2, 10 mM HEPES (pH 7), 0.5% Triton X-100, and 10mM VRC) for 7 min. Cells were washed in PBS and fixed in 4% formaldehyde for 10 min followed by consecutive dehydration with 70%, 85%, and 100% ethanol. Cells were then rehydrated in 2X SSC 50% formamide before hybridization with a Cy3-conjugated PNA-(CCCTAA)_3_ probe for 16 h. After hybridization, cells were washed with 2X SSC 50% formamide and directly used for immunofluorescence to visualize FLAG-TPP1 constructs. Imaging was performed as described above.

#### Detection of nascent telomeric synthesis using mutant TR (IF-coFISH)

For detection of newly synthesized mutant telomeric DNA in HeLa, FLAG-TPP1-S, and FLAG-TPP1-L M87A stable cell lines, cells were first induced with doxycycline for 48 h prior to transfection with mutant telomerase. After 48 h induction, 1 µg of pTERT-cDNA6/myc-His C and 3 µg of phTRmut-Bluescript II SK (+) plasmids were transfected into approximately 1 million cells with lipofectamine LTX (Fisher; 15338100) using the manufacturer recommended protocol. 20-24 h post transfection, immunofluorescence was performed exactly as described above to visualize FLAG-TPP1 constructs. Subsequently cells were analyzed by FISH to detect mutant TR and newly synthesized mutant telomeres. Cells were fixed with 4% formaldehyde for 10 min at room temperature, washed with PBS, and consecutively dehydrated with 70%, 95%, and 100% ethanol. Cells were then rehydrated with 2X SSC 50% formamide and pre-hybridized for 1 hour using hybridization solution containing 100mg/ml dextran sulfate, 0.125 mg/ml yeast tRNA, 1 mg/ml BSA, 0.5 mg/ml salmon sperm DNA, 1 mM vanadyl ribonucleoside complexes (VRC), and 50% formamide in 2X SSC. After pre-hybridization cells were transferred to hybridization solution containing a mixture of three Cy5-conjugated probes against TR (described previously) and a Cy3-conjugated PNA-(CCGCAA)_3_ probe. Cells were heat denatured at 80°C for 5 min then incubated at 37°C for ~16 h. After incubation cells were washed with 2X SSC 50% formamide and mounted on microscope slides using ProLong Gold mounting medium with DAPI (Life Technologies). Detection of newly synthesized mutant telomeres in HeLa cells transiently transfected with FLAG-TPP1-L M87A or FLAG-TPP1-L M87A R_3_E_3_ was performed similarly 40 h after transfection with 1 µg FLAG-TPP1 construct, 1 µg of pTERT-cDNA6/myc-His C plasmid, and 3 µg of phTRmut-Bluescript II SK (+) plasmid. HeLa cells with no exogenous TPP1 expression were analyzed both 20 h and 40 h after transfection with the TERT and mutant TR plasmids. All imaging was performed using a laser scanning confocal microscope (SP5; Leica, Germany) equipped with a 100X oil objective. ImageJ and Adobe Photoshop were used to process all images. Colocalization between TR and mutant telomere signal was quantified manually by two separate individuals. Mutant telomere signal was only scored when it was detectible above the background signal (lower pixel intensity threshold set at 30, upper pixel intensity threshold set at 255). Statistical significance was scored using a two-tailed student’s t test.

### Co-immunoprecipitation assay for protein-protein interaction

Co-immunoprecipitation experiments were performed as described previously (Grill et al., 2018). Briefly, HeLa-EM2-11ht cells were transfected with 1 µg each of plasmids containing Flag-TPP1-S, Flag-TPP1-M87A, Myc-POT1, FLAG-POT1, or Myc-TIN2. 48 h after transfection/ induction with doxycycline, cells were washed with PBS, trypsinized, and dislodged with medium containing 50% FBS. Cells were then pelleted, resuspended in 400 µL of lysis buffer (50 mM Tris-Cl (pH 7.6), 20% glycerol, 1 mM EDTA, 150 mM NaCl, 0.5% Triton X-100, 0.02% SDS, 1 mM dithiothreitol, 2 mM phenylmethylsulfonyl fluoride, complete protease inhibitor cocktail [Roch]) and kept on ice. 33 µL of 4 M NaCl, and 433 µL of water was added and lysates were centrifuged at 13,600 rpm for 10 min. After centrifugation, 40 µL of lysate was added to SDS gel loading buffer and kept aside for analysis of input samples. Remaining supernatant was used directly for immunoprecipitation with 30 µL of pre-washed anti-FLAG M2 affinity gel (Sigma; A2220). After overnight incubation, beads were washed and protein was eluted from the beads by adding 60 µL of 2X SDS gel loading buffer. All samples were analyzed by SDS-PAGE followed by immunoblotting with HRP conjugated FLAG or MYC antibodies.

### Immunoblotting

Immunoblotting was performed using standard procedures with the following antibodies: mouse monoclonal anti-FLAG M2-HRP conjugate (Sigma; A8592; 1:10,000), mouse monoclonal anti-c-Myc (9E10) HRP conjugate (Santa Cruz; sc-40 HRP; 1:10,000), mouse monoclonal anti-β-actin antibody (Sigma; A5441; 1:10,000), and rabbit polyclonal TPP1 antibody (Bethyl laboratories; A303-069A; 1:2,500). Secondary horseradish peroxidase-conjugated goat antibodies against rabbit or mouse IgG (Santa Cruz Biotechnology; 1:10,000) were used for detection with chemiluminescence by ECL plus reagents (Pierce ECL Western Blotting Substrate; Thermo Scientific). All antibodies used to assess knockdown of components in RNA decay and processing events were a kind gift from Dr. Peter Baumann. The rat monoclonal antibody against Ago2 was a kind gift from Dr. Gunter Meister. The rabbit polyclonal anti-Dicer antibody was a kind gift from Dr. Nils Walter. The data were visualized and quantified using a gel-documentation system (ChemiDoc™ MP System; BioRad).

### Testes tissue processing and immunoblotting

Normal testes tissue was obtained from the Tissue Core of the University of Michigan Rogel Cancer Center. Testes tissue used for immunoblotting was homogenized in cell extraction buffer (Invitrogen; FNN011) supplemented with PMSF and protease inhibitor cocktail (Sigma 11836170001 ROCHE). Homogenized tissue was incubated on ice for 1 h followed by centrifugation at 4°C and 13,000 rpm for 15 min. Soluble lysate was removed and the resulting pellet was resuspended in buffer containing 5 M urea, 2 mM MgCl2, 4% SDS, 140 mM Tris-Cl (pH 8), and 50 mM DTT. The resulting chromatin fraction was boiled with 5X SDS loading dye and analyzed by SDS-PAGE and immunoblotting with rabbit polyclonal TPP1 antibody following standard procedures.

### Enrichment of endogenous TPP1 using POT1-DNA pull down

Immunoprecipitation experiments to detect endogenous TPP1 protein were performed by first incubating 10 µg of purified POT1 protein with a biotinylated telomeric DNA primer (5’-biotin-ACGTA(GGTTAG)_3_) bound to 20 µl of streptavidin beads (Thermo; Pierce; 20357) for 2 h on ice. During this incubation, approximately 4 million HeLa, HEK 293T, and BJ fibroblast cells (kind gift from Dr. Ursula Jakob) were lysed in 50 mM Tris-Cl (pH 7.6), 20% glycerol, 1 mM EDTA, 150 mM NaCl, 0.5% Triton X-100, 0.02% SDS, and 1 mM DTT and centrifuged at 4°C and 13,000 rpm for 10 min. After 2 h, the soluble lysates were added to the POT1-DNA-bead slurry and incubated for ~16 h at 4°C. Beads were then washed and proteins were eluted with 2X SDS loading buffer. Samples were analyzed by SDS-PAGE and immunoblotting with TPP1 antibody.

### CRISPR-Cas9 mediated insertion of a C-terminal FLAG tag at the endogenous locus of human TPP1

Insertion of the FLAG tag into the HEK 293T genome was performed using a strategy reported previously for introducing the K170Δ into the HEK 293T genome (Bisht et al., 2016). The design of the ssODN is shown in Figure S4B and involves insertion of a 3X-FLAG coding DNA sequence in-frame and immediately upstream of the natural stop codon of TPP1-L and TPP1-S. Single clones were analyzed using FLAG-immunoblotting.

### CRISPR-Cas9 mediated TPP1-L KO generation

Generation of TPP1 knockout clones was performed using a strategy previously reported (Bisht et al., 2016). A guide RNA was designed with the Zhang Lab open access CRISPR design tool. The ssODN (described in Fig. 4E) was designed to encompass the desired mutations as well as a silent *Aat*II restriction endonuclease cleavage site. A surveyor nuclease assay (Surveyor® Mutation Detection Kit; catalog #706025; Transgenomic) was used to determine cleavage efficiency 48 h after transfection of 1 µg of guide RNA and Cas9 encoding plasmids into HeLa-EM2-11ht cells following the manufacturer’s protocol. Upon completion of the reaction, products were visualized using ethidium bromide-stained agarose gels and robust cleavage was observed. HeLa-EM2-11ht cells were then transfected with guide RNA and Cas9 encoding plasmids along with indicated ssODN as previously described. 3 d after transfection 2,000 cells were plated for colony formation while the rest were used to test the efficiency of repair. To check repair efficiency genomic DNA was prepared using GenElute Mammalian Genomic DNA Purification Kit (#G1N70; Sigma) and the locus encompassing our desired mutations was PCR amplified. Amplified DNA was then digested in the presence or absence of *Aat*II and reaction products were visualized on an ethidium bromide stained agarose gel. ~2 weeks after seeding, colonies were picked and transferred to a 96 well plate where they remained for 1 week. They were then trypsinized and half the cells were transferred to a replicate plate for genomic DNA preparation. Isolated genomic DNA from each clone within the 96 well plate was PCR amplified and subjected to digestion by *Aat*II. Positive clones were expanded and genotyped using Sanger DNA sequencing. Selected clones were then propagated for telomere length analysis.

### 5’ RACE to clone sunRNA

5’ RACE was performed using the 5’ RACE kit from Life Technologies (#18374058). First strand cDNA synthesis was performed with SuperScript™ II RT using 2 μg total RNA from HEK 293T cells, a gene-specific antisense oligonucleotide GSP1 (TCCCTGATCCTCTCCTCTCC) and abridged anchor primer, AAP (GGCCACGCGTCGACTAGTACGGGIIGGGIIGGGIIG) which contains a complementary homopolymeric tail annealing sequence and allows amplification from a homopolymeric tail. Following cDNA synthesis, the first strand product was treated with RNAse and purified from unincorporated dNTPs and GSP1 using S.N.A.P. columns provided with the kit. 10 µl of S.N.A.P. column purified cDNA was tailed at the 3’ end with CTP using TdT (Terminal deoxynucleotidyl transferase) enzyme. After heat inactivation of TdT, 5 µl of tailed cDNA was amplified by PCR for 30 to 35 cycles using a nested gene-specific primer GSP2 (TCCTCTCCTCTCCTGCCGC) which anneals 3’ to GSP1 and a complementary homopolymer-containing abridged anchor primer (AAP) which permits amplification from the homopolymeric tail. 0.1% of the PCR reaction from this step was subjected to re-amplification using a nested gene specific primer GSP3 (GGAAGCAGAGTGTGGAGCGG) and abridged universal amplification primer AUAP (GGCCACGCGTCGACTAGTAC). The PCR products were resolved on a 2% GTG agarose gel and discrete bands were gel eluted and subjected to TA cloning and sequencing. Sequences showing a homopolymer C tail after sequencing were considered positive. All sequenced cDNA were positive for the 3’-UTR sequence of TPP1, but contained different extents of upstream exonic sequences.

### Cloning, expression, and silencing experiments with sunRNA

Protein-coding inserts contained cDNA for only the ORF cloned into multiple cloning site 1 of the bi-directional dox-inducible pTet-BI4 vector using *Not*1/*Xho*1 restriction based cloning. FLAG-TPP1-L was cloned into pTet-Bi4 vector with a 3X-FLAG tag on the C-terminus and a 1X-FLAG tag on N-terminus of the TPP1-L protein and used in all sunRNA experiments requiring TPP1-L expression. The sunRNA sequence was similarly cloned into multiple cloning site 2 using *Spe*1/*Nhe*1 sites. Because *Spe*1 and *Nhe*1 generate compatible sticky ends, the same inserts, ligation reactions, and transformations were used to generate plasmid constructs with sunRNAs and cognate reverse complements (RC). SV40 polyA sequences downstream of the multiple cloning sites were used for proper RNA maturation of the cloned protein and sunRNA/RC cDNA constructs. Plasmids encoding TPP1-L (or other protein-coding genes used in the study) and Vector/sunRNA/RC cDNA were transfected into HeLa-EM2-11ht as already described. 24 h post-transfection, cells were lysed in SDS-gel loading buffer and analyzed by immunoblotting (for detecting proteins) as already described, or by Northern blot analysis as described below.

### Northern blotting

10 μg of total RNA was dissolved in buffer containing 5.7 μl DEPC-treated water, 1 μl 10 X MOPS, 3.3 μl formaldehyde, and 10 μl formamide and mixed well. The mixture was heated at 72°C for 5 min, followed by the addition of 5 μl of 6X DNA loading dye and transfer to ice. The agarose gel for running the RNA samples was prepared as follows. 126.75 ml water was added to 1.5 g agarose heated in a microwave to get a homogenous solution that was kept aside until the temperature reduced to approximately 55 **°**C. 10 ml 10X MOPS, 8.25 ml 37% formaldehyde, and 3 μl ethidium bromide were heated in a separate tube in a 65°C water bath. The two solutions were mixed together and cooled to form the gel matrix for electrophoresis. The RNA samples were loaded on the gel and run in 1X MOPS buffer at 50 V until the bromophenol blue front traversed approximately half the length of the gel. The gel was visualized under UV light to detect and capture an image of the bands for the 18S and 28S ribosomal RNA subunits. The gel was then transferred to Hybond nylon membrane using 10X SSC overnight. The protocol described for Southern blotting was applied for detecting the indicated RNA bands in the figures. A mixture of five probes, TPP1 ORF: 1R, 2R, 3R, 4R, and 5R, were used for detecting the TPP1-L or TPP1-S mRNA. All these probes were complementary to regions common to both TPP1-L and TPP1-S mRNA. Probes containing the TPP1 3’-UTR sequence and its reverse complement were used to detect RC and sunRNA, respectively.

### siRNA knockdown experiments

HeLa-EM2-11ht cells were transfected with 80 nmol of the indicated siRNA using Oligofectamine (Life Technologies) in a 24-well culture plate. All siRNA were a kind gift from Dr. Peter Baumann. 24 h after transfection, cells were trypsinized and split into three wells of a 12-well plate. 48 h post transfection, 500 ng plasmids encoding TPP1-L and vector/sunRNA/RC constructs were transfected using Lipofectamine 2000 (Life Technologies). 10 h after the second round of transfection, the media was changed to include doxycycline (200 ng/ml). Lysates were prepared 24 h post-induction and immunoblotting was performed as already described.

### Protein purification

TPP1^90-334^ and POT1 proteins were purified exactly as described previously (Grill et al., 2018; Nandakumar et al., 2012). TPP1^1-334^ was purified in parallel with TPP1^90-334^ using the same protocol.

### DNA-binding experiments

Filter-binding assays with POT1-DNA, TPP1^90-334^-POT1-DNA, and TPP1^1-334^-POT1-DNA complexes were performed and analyzed exactly as previously described (Nandakumar et al., 2012; Nandakumar et al., 2010).

### Direct telomerase activity assays

Telomerase assays were performed as described previously (Grill et al., 2018). Briefly, HEK 293T cells were transfected with 1 µg of pTERT-cDNA6/myc-His C and 3 µg of phTR-Bluescript II SK (+) plasmids and soluble cell extracts were prepared as previously described(Kocak et al., 2014). This extract was used as the source of telomerase enzyme in direct telomerase primer extension assays based on previously published protocols (Kocak et al., 2014). In each reaction, 2 µl of cell extract containing telomerase enzyme was incubated at 30°C for 10, 20, 30, or 60 min. Full reactions contained 50 mM Tris-Cl (pH 8.0), 1 mM MgCl2, 1 mM spermidine, 30 mM KCl, 5 mM β-mercaptoethanol, 1 µM of primer a5 (TTAGGGTTAGCGTTAGGG), 500 µM dATP, 500 µM dTTP, 2.92 µM unlabelled dGTP, 0.17 µM radiolabeled dGTP (3000 Ci/mmol). Purified POT1, TPP1^90-334^, or TPP1^1-334^ proteins were added to the reactions at indicated concentrations. 100 µl of buffer containing 3.6 M ammonium acetate and 20 µg of glycogen was used to quench each reaction, and the DNA products were precipitated with 70% ethanol. The pellets were resuspended in 7 µl water, then mixed with 7 µl loading buffer containing 95% formamide, and heated at 95°C for 5 min. Samples were resolved on a 10% acrylamide, 7M urea, 1X TBE sequencing-size gel and gels were dried then imaged on a phosphorimager (Storm; GE). All assays were analyzed using Imagequant TL (GE Life Sciences) software. Processivity calculations were performed as described previously (Grill et al., 2018).

### Visualization of data on UCSC genome browser

Indicated tracks were loaded on to the UCSC genome browser (Casper et al., 2018) and saved as. ps files. The sessions were processed in Adobe Illustrator to furnish final figures. All UCSC genome browser associated data were previously deposited in databases by other groups and have been cited appropriately.

### Testes-specific next-generation sequencing analysis of data deposited in the SRA (Sequence Read Archives)

Sequence reads accession files corresponding to each run were accessed from the Sequence Read Archives, downloaded, and converted to raw sequence data (.fastq files) using the prefetch and fastq-dump commands from the sra-tools v2.8.2 suite. Deduplication, quality-trimmming, and adapter removal was performed on all data sets using dedupe.sh, clumpify.sh, and bbduk.sh from the BBTools suite v37.90. Quality assessments were performed on all raw sequencing and processed data using FastQC v0.11.7 and MultiQC v1.5. Quality control processed reads were aligned to the human genome (UCSC hg19) (Casper et al., 2018), using hisat2 and the --dta option with all other defaults. Uniquely mapping reads or read-pairs were selected used for further analysis (See command below).

hisat2 --dta --new-summary -x {params.fasta_basename} -1 {input.R1} -2 {input.R2} -p {threads} 2> {output.summary} | perl -lane ’if($_ !~ m/NH:i:1/){{print}}’ | samtools view -@ {threads} -Sb - > {output.bam}

Aligned reads in binary alignment format (.bam files) were sorted and indexed using the samtools sort and samtools index commands from samtools v1.8. Read depth-normalized bigwig files were created from sorted alignment files for ease of viewing in genome browsers using the bamCoverage command from deepTools v3.0.2 with --binSize 10 and – normalizeUsing BPM options set. With the exception of the use of sra-tools, the workflow as implemented in Snakemake v4.7.0.

Programs/Softwares used:

SRA-Tools v2.8.2 (Leinonen et al., 2011)

FastQC v0.11.7 https://www.bioinformatics.babraham.ac.uk/projects/fastqc/

MultiQC v1.5 (Ewels et al., 2016)

hisat2 v2.1.0 (Kim et al., 2015)

samtools v1.8 (Li et al., 2009)

perl v5.22.0.1

deeptools v3.0.2 (Ramirez et al., 2016)

BBTools v37.90 https://jgi.doe.gov/data-and-tools/bbtools/

Snakemake v4.7.0 (Koster and Rahmann, 2018)

## Supplemental Figure legends

**Supplemental Fig 1.**
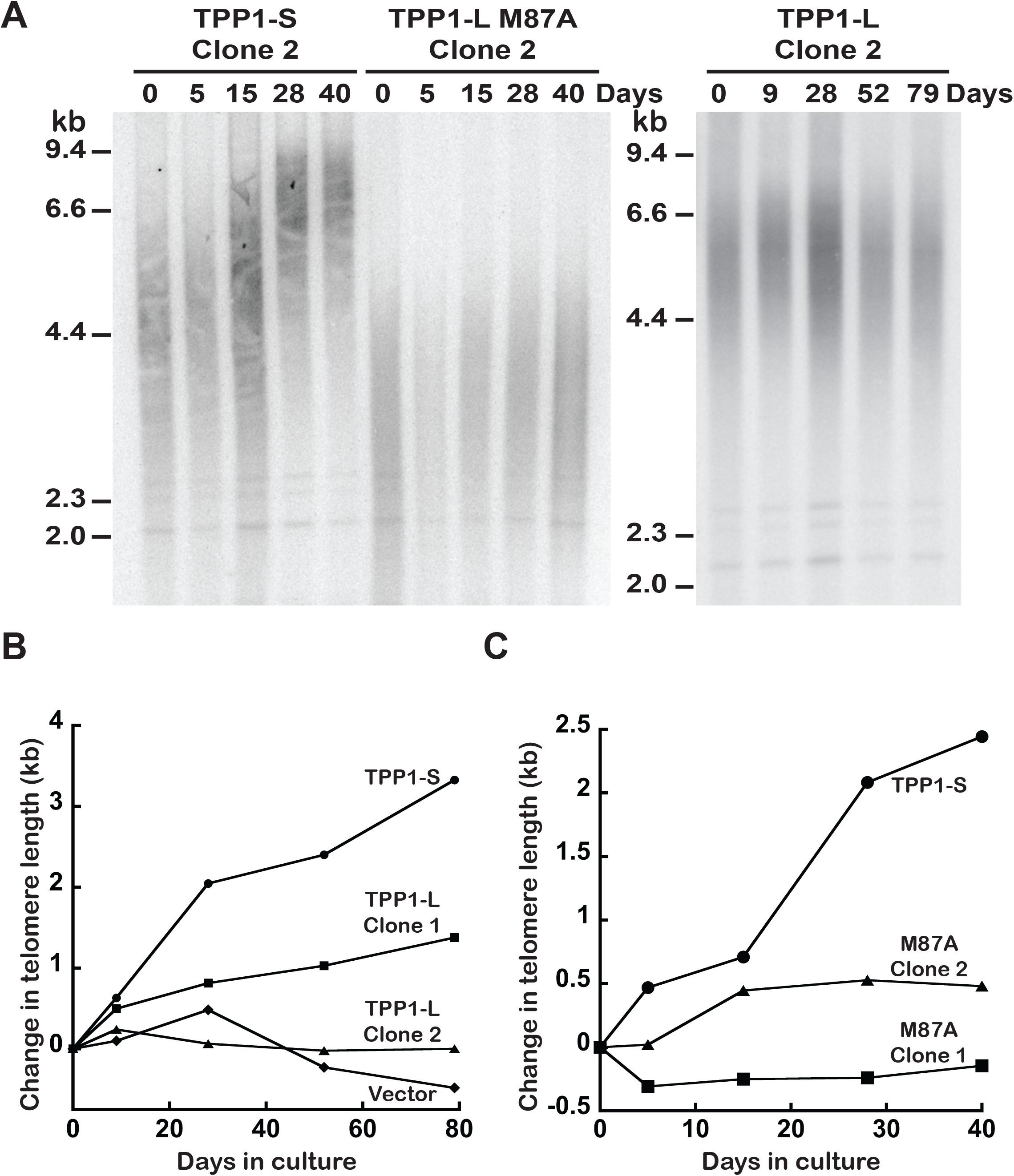
Telomere restriction fragment length analysis of TPP1-L and TPP1-S clonal cell lines. Related to Figure 1. (A) Telomere restriction fragment (TRF) analysis of a second clone for HeLa-EM2-11ht cells expressing TPP1-S, TPP1-L, or TPP1-L M87A constructs for the indicated number of days in culture. (B and C) Plot of the change in mean telomere length for HeLa-EM2-11ht cells stably expressing the indicated constructs for the indicated number of days in culture.

**Supplemental Fig. 2.**
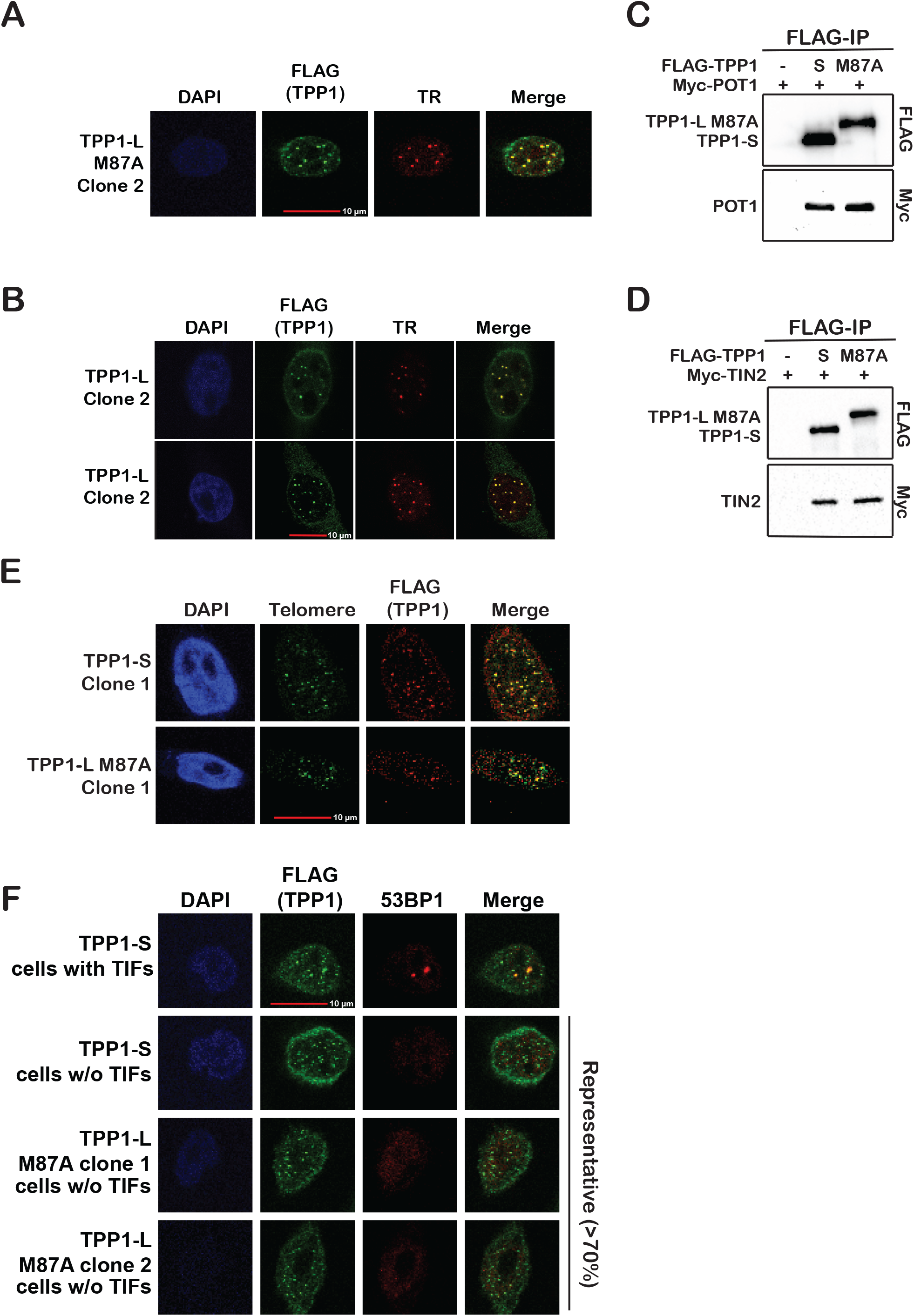
Both TPP1-L and TPP1-S are proficient in chromosome end protection and telomerase recruitment. Related to Figure 2. (A and B) IF/FISH analysis of telomerase recruitment for TPP1-L M87A clone #2 (A) and two clones of TPP1-L WT (B). “FLAG (TPP1)” refers to immunofluorescence signal of FLAG-TPP1-L or FLAG-TPP1-L M87A at telomeres (green). “TR” represents FISH signal for telomerase RNA (red). “Merge” demonstrates successful telomerase recruitment to telomeres (yellow). See also Figure 2 for quantitation. (C) Pull down of transiently expressed FLAG-TPP1-S or FLAG-TPP1-L M87A and Myc-POT1 on anti-FLAG conjugated beads. (D) Pull down of transiently expressed FLAG-TPP1-S or FLAG-TPP1-L M87A and Myc-TIN2 on anti-FLAG conjugated beads. (E) IF-FISH depicting colocalization of indicated TPP1 construct with telomeres. “Telomere” refers to signal from a C-rich, Cy3-labled PNA probe against telomeric DNA (green). “FLAG (TPP1)” refers to FLAG immunofluorescence signal of indicated TPP1 construct (red). “Merge” panel reveals extent of TPP1 colocalization to the telomere (yellow). (F) Cells stably expressing TPP1-S or TPP1-L M87A were analyzed for TIFs using co-immunofluorescence (co-IF) against “FLAG (TPP1)” as a telomere marker and “53BP1” as a DNA damage marker. Co-localization of 53BP1 foci and FLAG-TPP1 foci within “Merge” panels indicates DNA damage at the telomere. An example of a cell with TIFs is shown along with images of cells that are more representative of the majority of cells.

**Supplemental Fig. 3.**
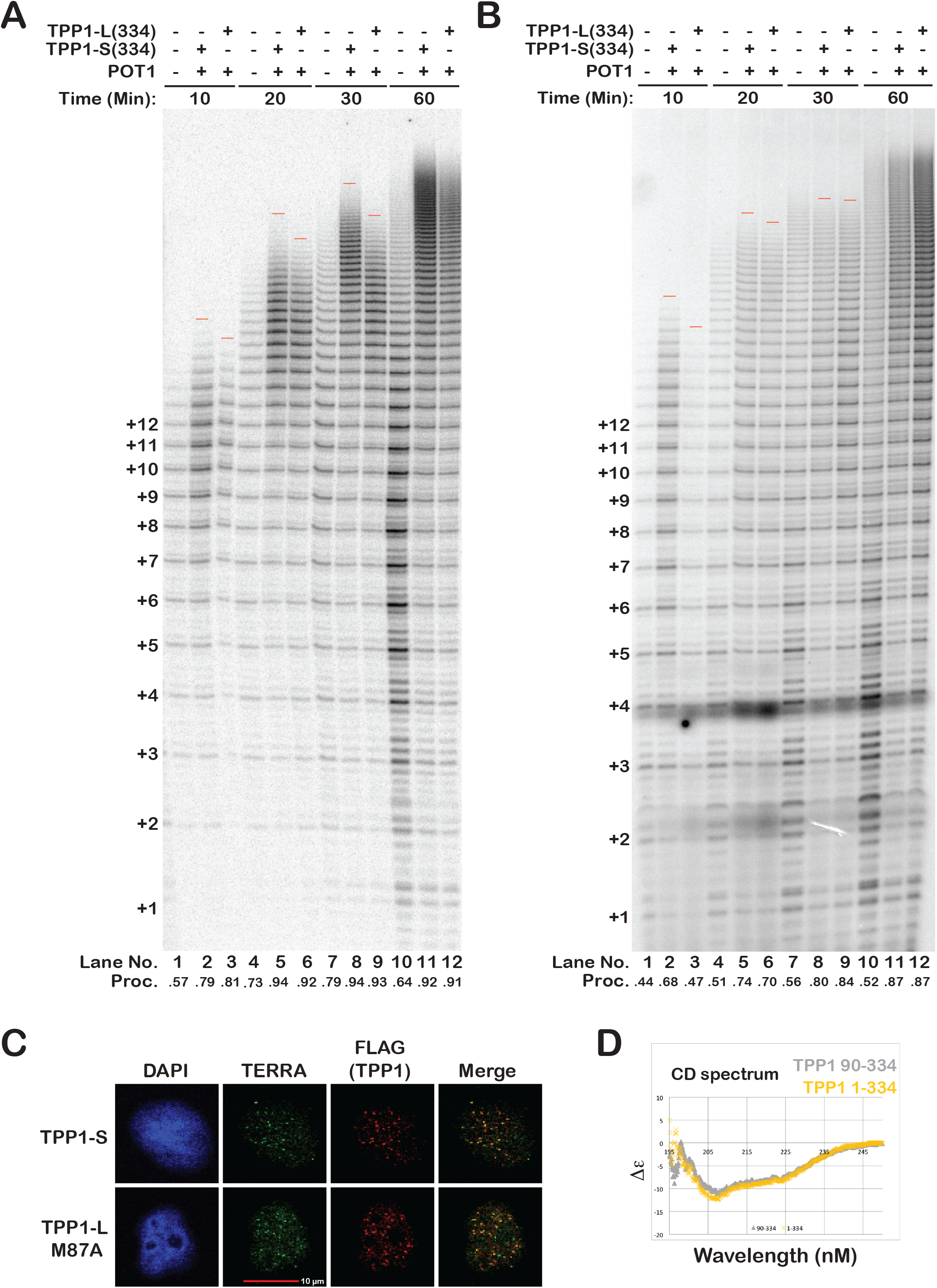
Additional replicates of direct telomerase activity assays with TPP1-L and TPP1-S proteins. Related to Figure 3. (A and B) Direct telomerase activity assay with purified POT1, TPP1-S^90-334^, and TPP1-L^1-334^ proteins, as shown in Figure 3. (C) IF/RNA-FISH analysis of TERRA colocalization with the indicated TPP1 construct. “TERRA” depicts signal from a C-rich, Cy3-labled telomeric PNA probe bound to RNA. “FLAG (TPP1)” depicts immunofluorescence signal from indicated TPP1 construct. “Merge” represents extent of TERRA co-localization with the telomere (yellow). (D) Circular Dichroism (CD) spectra of TPP1^1-334^ and TPP1^90-334^ proteins showing almost superimposable profiles and suggesting little secondary structure in TPP1-L aa 1-86.

**Supplemental Fig. 4.**
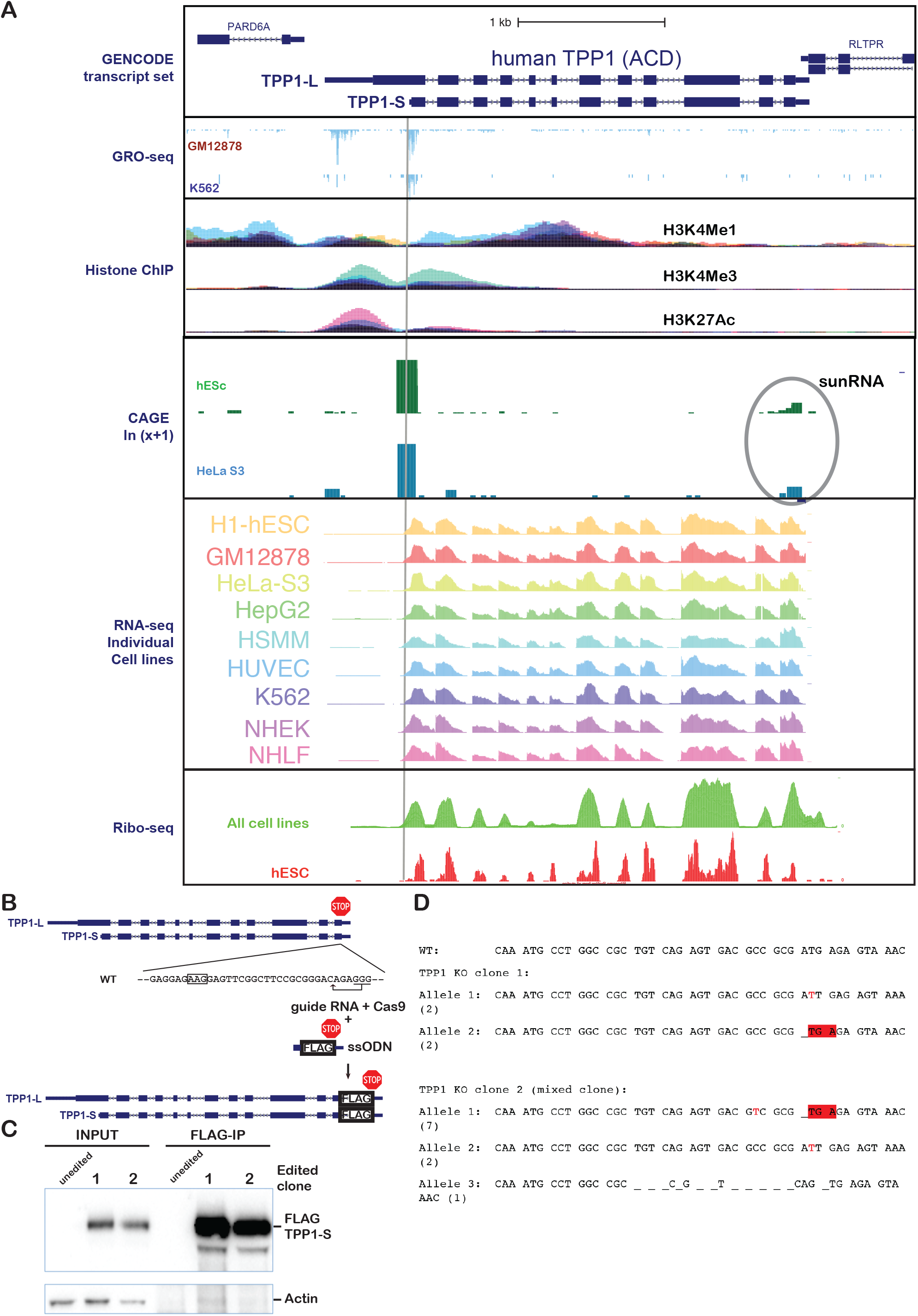
Transcriptome and protein analysis suggest separate transcription cassettes for TPP1-L and TPP1-S, although TPP1-S protein is the predominant form in cells. Related to Figure 4. (A) High-throughput sequencing data visualized on the UCSC genome browser at the *ACD* locus. GRO-seq data for assessing nascent mRNA from the indicated cell lines, cumulative ChiP data from a collection of cell lines for indicated histone modifications for assessing gene expression potential, CAGE data for human embryonic stem cells and HeLa S3 cells, RNA-seq tracks from individual cell lines, and Ribo-seq data (cumulative versus embryonic stem cell line) are shown along with the TPP1-L and TPP1-S transcript annotations from GENCODE. The vertical grey bar denotes the 5’ end of TPP1-S mRNA. (B) CRISPR-Cas9 strategy to introduce the 3X FLAG tag at the C-terminus of endogenous TPP1 in HEK 293T cells is shown. (C) Immunoprecipitation of lysates from clones isolated from the FLAG-TPP1-S CRISPR experiment using anti-FLAG M2 affinity gel followed by blotting using the anti-actin and anti-FLAG-HRP conjugate. An unedited clone was included as negative control. (D) Allele status of *ACD* (TPP1) in the CRISPR-mediated KO of TPP1-L.

**Supplemental Fig. 5.**
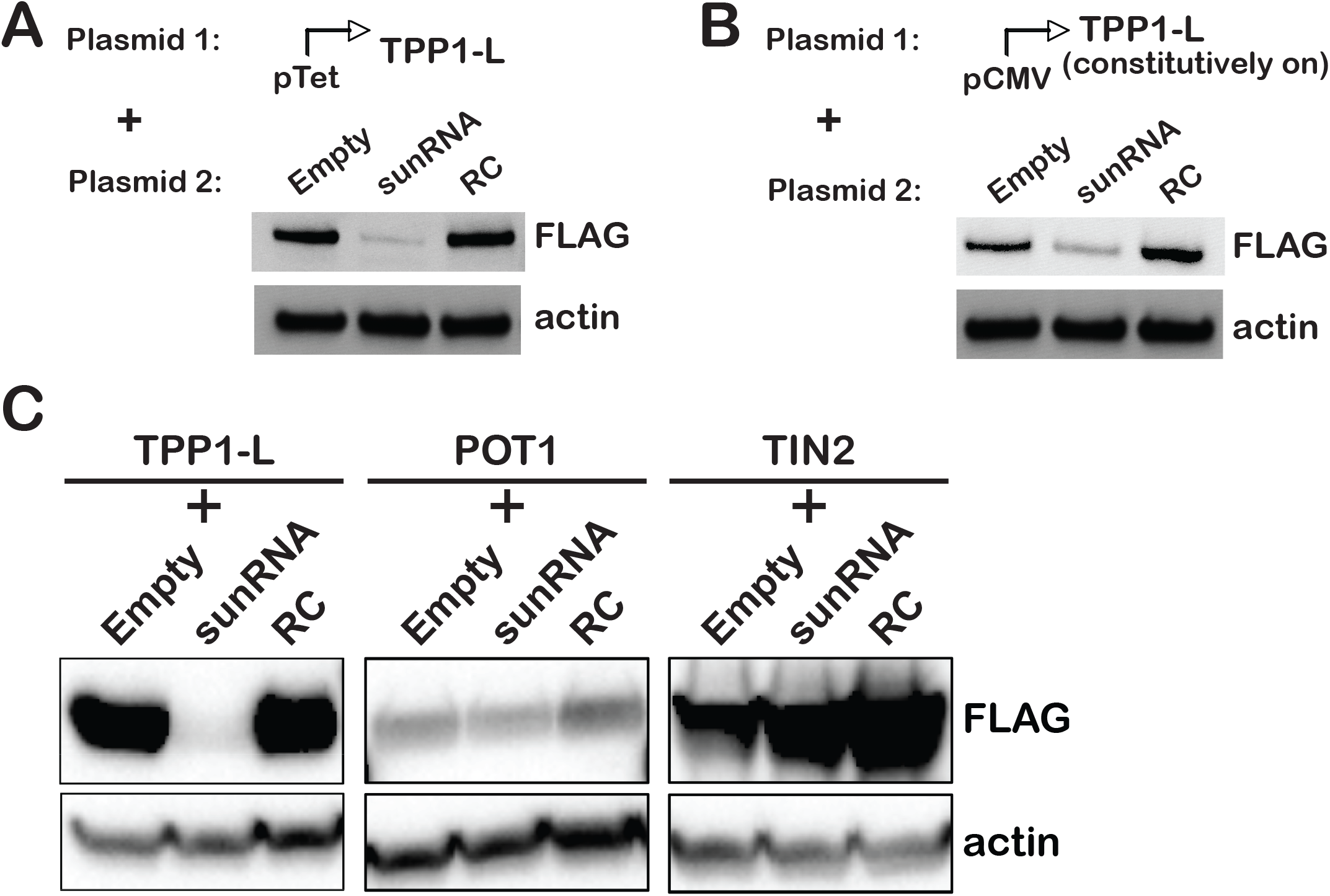
Silencing of the sunRNA is specific for TPP1-L and independent of the vector context. Related to Figure 5. (A) Anti-FLAG immunoblots were performed on lysates from cells co-transfected with two plasmids, one of which codes for dox-inducible FLAG-TPP1-L, and the other for dox-inducible empty, sunRNA, or RC constructs. (B) Anti-FLAG immunoblots were performed on lysates from cells co-transfected with two plasmids, one of which codes for FLAG-TPP1-L driven by a constitutive CMV promoter, and the other for dox-inducible empty, sunRNA, or RC constructs. (C) Anti-FLAG immunoblots showing that the sunRNA of TPP1 silences TPP1-L but not human POT1 or TIN2.

**Supplemental Fig. 6.**
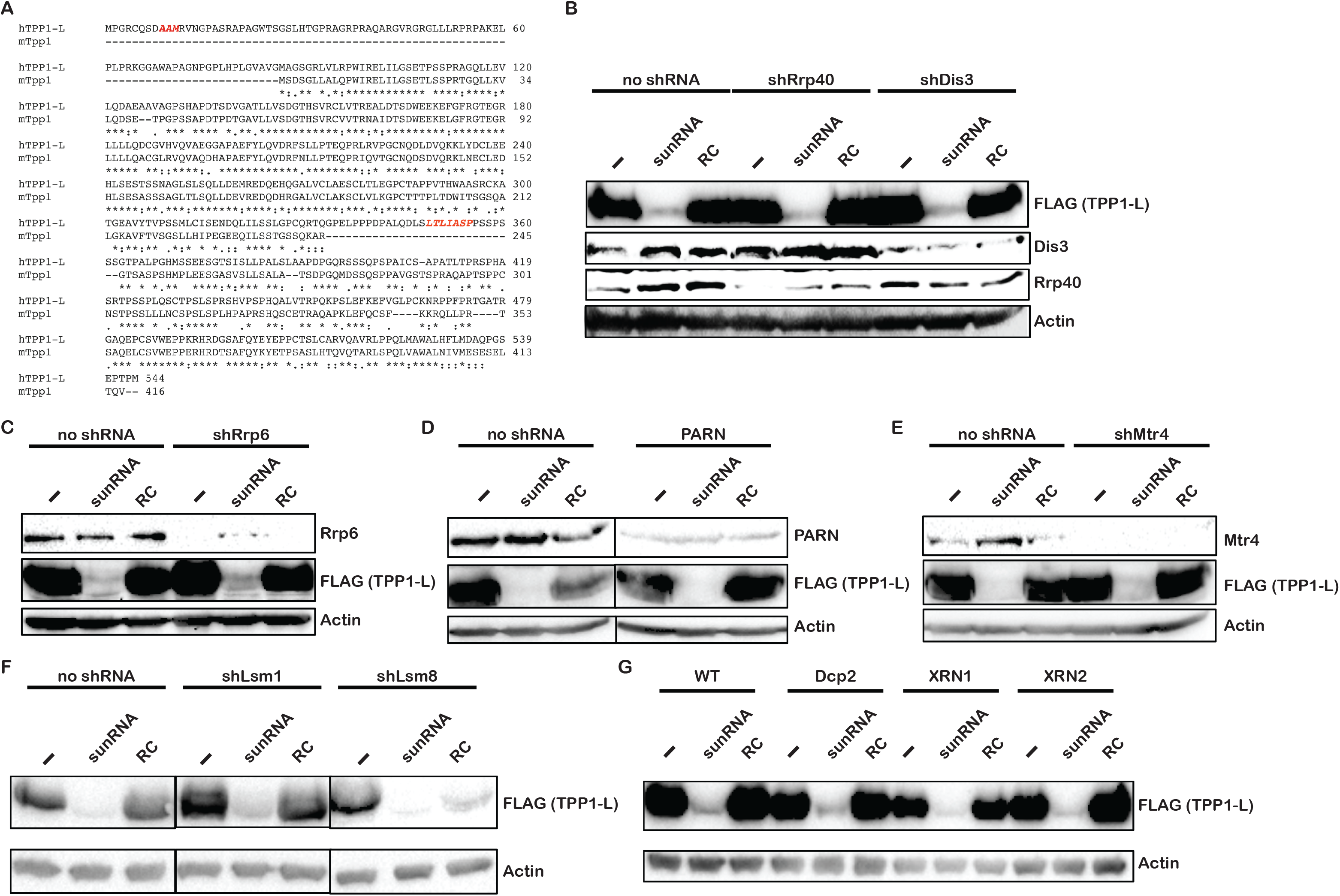
sunRNA mediated regulation of TPP1 is specific to human TPP1 sequence but not dependent on any unique RNA surveillance or silencing machinery. Related to Figure 6. (A) Amino acid sequence alignment of human TPP1-L and mouse TPP1. Residues indicated in red are coded by nucleotides predicted to be complementary to the human sunRNA sequence. Both these motifs reside in regions that are absent in mouse TPP1 protein (and mRNA). (B-G) sunRNA-mediated silencing of TPP1-L prevails despite siRNA-mediated knockdown of Rrp40, Dis3 (B); Rrp6 (C); PARN (D); Mtr4 (E); LSM1, LSM8 (F); Dcp2, XRN1, or XRN2 (G). “-“ indicates Empty vector control.

